# Identification of Gip as a novel phage-encoded gyrase inhibitor protein featuring a broad activity profile

**DOI:** 10.1101/2021.01.28.428610

**Authors:** Larissa Kever, Max Hünnefeld, Jannis Brehm, Ralf Heermann, Julia Frunzke

## Abstract

Bacteriophages represent a powerful source for the identification of novel antimicrobial proteins. In this study, a screening of small cytoplasmic proteins encoded by the CGP3 prophage of *Corynebacterium glutamicum*, resulted in the identification of the novel <u>g</u>yrase-<u>i</u>nhibiting <u>p</u>rotein Cg1978 (Gip), which shows a direct interaction with the gyrase subunit A (GyrA). In vitro supercoiling assays further suggest a stabilization of the cleavage complex by Gip. Overproduction of Gip in *C. glutamicum* resulted in a severe growth defect as well as an induction of the SOS response. The cells adapted to *gip* overexpression by increasing expression levels of *gyrAB* and by reducing *topA* expression reflecting the homeostatic control of DNA topology. Interestingly, Gip features a similar activity profile towards gyrases of *Escherichia coli, Mycobacterium tuberculosis* and *C. glutamicum*. Therefore, the detailed elucidation of the mechanism of Gip action may provide novel directions for the design of drugs targeting DNA gyrase.

## Introduction

Bacteriophages represent the ‘dark matter’ of the biological world (Hatfull, 2015; Ofir & Sorek, 2018; Rohwer & Youle, 2011). With the recent massive expansion in the genomic sequence space, also the number of functionally unknown open reading frames (ORFans) in phage genomes is continuously increasing (Yin & Fischer, 2008). By targeting diverse cellular processes and regulatory hubs in their host cell, bacteriophages represent a rich source for the identification of novel antibacterial proteins as well as for the establishment of highly efficient molecular tools (De Smet et al., 2017; Nobrega et al., 2018; Roach & Donovan, 2015; Schroven et al., 2021). Especially, small cytoplasmic phage proteins have been shown to influence and reprogram a variety of key cellular processes, including transcription, translation, cell division or central metabolism (De Smet et al., 2017; Orr et al., 2020; Storz et al., 2014).

The DNA gyrase represents a type IIA topoisomerase present in all bacteria and plays a crucial role in the homeostatic control of DNA topology. Because of its unique ability to introduce negative supercoils into covalently linked double-stranded DNA, gyrase is an essential protein for bacterial cells and a key target of antibacterial agents (Bush et al., 2015). The heterotetrameric enzyme consists of two GyrA and two GyrB subunits (GyrA2GyrB2). While the GyrA subunit of the enzyme catalyzes the breaking and resealing activity, the GyrB subunit harbors the site of ATP hydrolysis (Reece & Maxwell, 1991; Vanden Broeck et al., 2019). In terms of small molecules, there are currently two major classes of drugs targeting the bacterial gyrase: the aminocoumarins and the quinolones (Bush et al., 2015; Collin et al., 2011). Besides a range of small molecules, also some proteinaceous bacterial toxins were found to inhibit the activity of the gyrase, including microcin B17 (Pierrat & Maxwell, 2003), a glycine-rich peptide found in *E. coli* strains carrying the *mcb* operon, as well as the CcdB toxin as part of the *ccd* toxin-antitoxin system encoded by the F-plasmid (Dao-Thi et al., 2005; Miki et al., 1992).

*Corynebacterium glutamicum* – a member of the phylum Actinobacteria – is an important industrial platform organism used for the industrial production of a wide range of value-added compounds, including amino acids, organic acids and proteins (Wendisch et al., 2016). The genome of the model organism *Corynebacterium glutamicum* ATCC 13032 contains four cryptic prophages (CGP1-4) (Frunzke et al., 2008; Ikeda & Nakagawa, 2003). The largest prophage CGP3 (~ 219 kb, including also CGP4) was shown to be inducible in an SOS-dependent as well as in an SOS-independent manner (Helfrich et al., 2015; Nanda et al., 2014; Pfeifer et al., 2016). Recently, the Lsr2-type protein CgpS was identified as a prophage-encoded nucleoid-associated protein involved in the silencing of phage gene expression maintaining the lysogenic state of the large CGP3 prophage (Pfeifer et al., 2016). Interference with CgpS binding was shown to result in prophage activation and consequently cell death.

In this study, a systematic screening of small cytoplasmic proteins encoded by the CGP3 prophage of *C. glutamicum* resulted in the identification of phage proteins causing severe growth defects and triggering prophage induction. The small phagic protein Cg1978 was further shown to directly target the DNA gyrase enzyme by interacting with the GyrA subunit. Interaction of Gip and GyrA was shown to inhibit the supercoiling activity of the DNA gyrase in vitro. Cg1978 was therefore termed Gip for <u>g</u>yrase <u>i</u>nhibiting <u>p</u>rotein. The function of Gip as a potential phage-encoded gyrase inhibitor is further supported by transcriptome analysis showing the upregulation of *gyrA* and *gyrB* transcripts as a compensatory mechanism to overcome Gip inhibition. Although Gip was shown not to be essential for prophage induction, we hypothesize that Gip plays a role in modulating gyrase activity to enable efficient phage DNA replication.

## Results

### Screening of small CGP3-encoded proteins for an impact on growth and prophage induction

Most of the proteins encoded by the cryptic but still inducible CGP3 prophage are of unknown function. Phage proteins were shown to frequently target key regulatory hubs to shut down bacterial metabolism (Roach & Donovan, 2015). In this study, we screened the impact of overall eleven small (<75 amino acids), cytoplasmic phage-encoded proteins on cellular growth and prophage induction. For this purpose, plasmid-based overexpression (pAN6-*GOI*) of the selected genes of interest was performed in the prophage reporter strain *C. glutamicum* ATCC 13032::P*_lys_*-*eyfp*. In previous studies, this genomically integrated reporter (*P_lys_*-*eyfp*) was successfully established to translate prophage activation into a fluorescence output (Helfrich et al., 2015). Following this approach, the overproduction of nine out of eleven phage proteins (Cg1902, Cg1910, Cg1924, Cg1925, Cg1971, Cg2026, Cg2035, Cg2045, Cg2046) displayed comparable phenotypes as the empty vector control regarding backscatter signal and fluorescence output measured via flow cytometry (Supplementary Figure S1 A and B). By contrast, overproduction of Cg1914 and Cg1978 showed a significant effect on growth and prophage induction. Cg1914 overproduction resulted in a reduced growth rate (μ = 0.26 ± 0.03 h^−1^) and a reduced final backscatter (Figure 1A, *blue line*) compared to the empty vector control pAN6 (μ = 0.40 ± 0.03 h^−1^). In case of Cg1978, overproduction led to an elongated lag-phase, but the final backscatter as well as the growth rate in the exponential phase (μ = 0.38 ± 0.01 h^−1^) (Figure 1A, *red line*) were comparable to those of the negative control.

**Figure 1:**
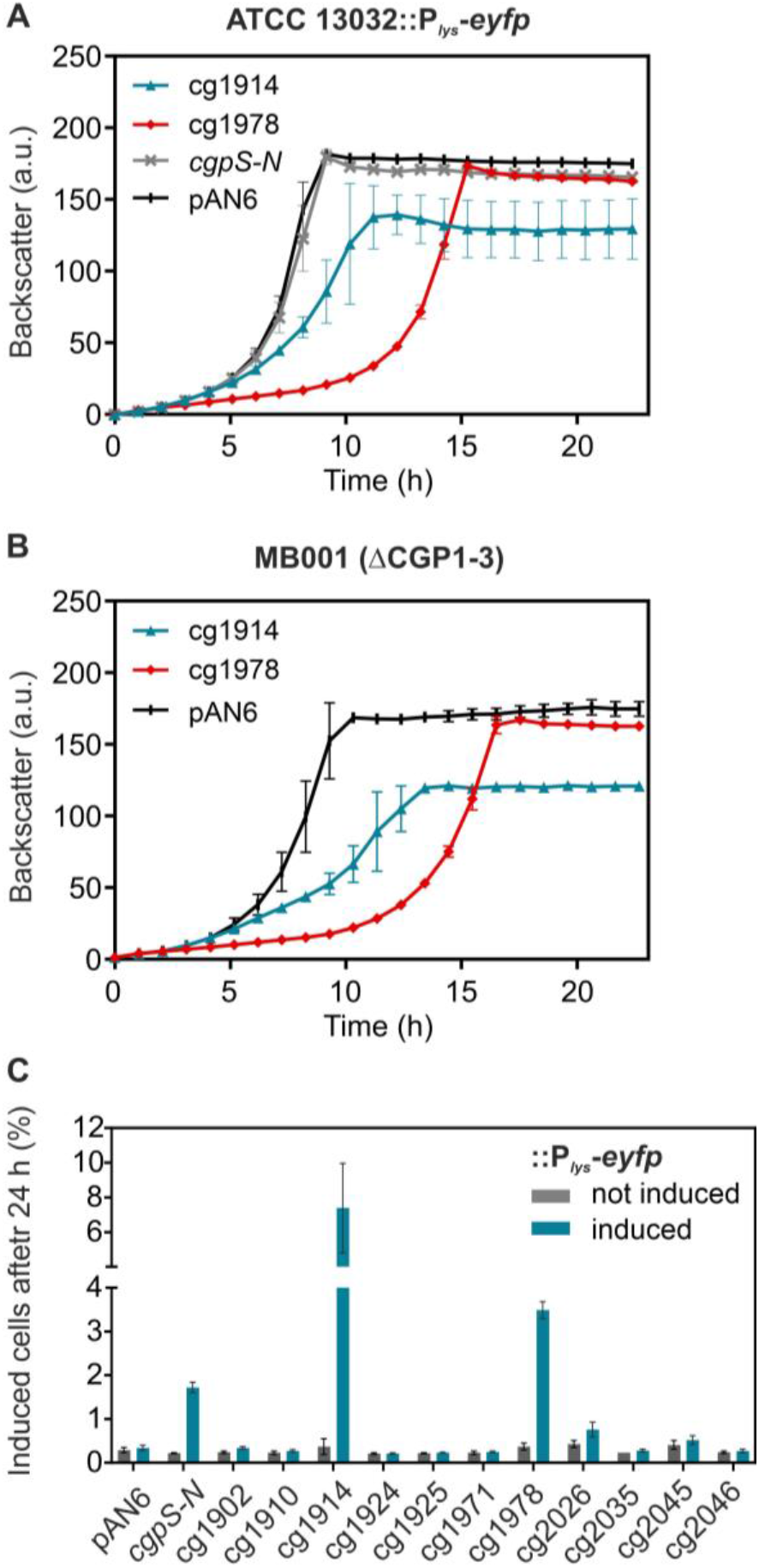
Screening of small phagic proteins regarding their impact on cellular growth and CGP3 induction in *C. glutamicum*. The cultivation of the prophage-reporter strain *C. glutamicum* ATCC 13032::P_*lys*_-*eyfp* and the prophage-free strain MB001 carrying the corresponding gene sequences of the small proteins on the pAN6 vector (under control of P*_tac_*) was performed in CGXII-Kan_25_ medium with 2 % (w/v) glucose and 50 μM IPTG for 24 h. All data represent mean values with standard deviations from three independent biological triplicates (n=3). **(A)** Growth curves of the prophage reporter strain (*C. glutamicum* ATCC 13032::P*_lys_*-*eyfp*) upon small protein overproduction are based on the backscatter measurements in the BioLector^®^ microcultivation system. **(B)** Growth curves of the prophage-free strain MB001 upon small protein overproduction are based on the backscatter measurements in the BioLector^®^ microcultivation system. **(C)** Percentage of induced cells after 24 h cultivation without and with 50 μM IPTG based on the flow cytometric measurements of the prophage reporter strain *C. glutamicum* ATCC 13032::P*_lys_*-*eyfp*.

A comparable impact on cell growth due to cg1914 or cg1978 overexpression was also detected in the prophage-free strain MB001, indicating that the observed growth defect was independent of the presence and/or activity of the CGP3 island (Figure 1B).

For both target proteins, Cg1914 and Cg1978, overproduction resulted in an increased fluorescence signal after 24 h cultivation (cg1914: 7.4 ± 2.6 % induced cells, cg1978: 3.5 ± 0.2 % induced cells) indicating CGP3 prophage induction in the respective subpopulation (Figure 1C). As a positive control, we expressed an N-terminally truncated variant of the prophage silencer CgpS (CgpS-N), which was previously shown to trigger prophage induction (Pfeifer et al., 2016). Since overproduction of Cg1914 and Cg1978 showed a high impact on prophage induction, we tested inducibility of the CGP3 prophage in mutants lacking the respective genes using a plasmid-based prophage reporter (P_*lys*_-*venus*). Remarkably, the corresponding strains *C. glutamicum* ATCC 13032 Δcg1914 and Δcg1978 featured no difference - neither in cell growth nor in prophage inducibility - upon treatment with the DNA-damaging antibiotic mitomycin C (Tomasz, 1995), which was used to trigger SOS-dependent prophage induction (Supplementary Figure S2). These results indicated that both proteins are not essentially involved in SOS-dependent CGP3 induction. Prophage induction, therefore, appeared to be an indirect effect of Cg1914 or Cg1978 overproduction. Based on further results described in the following, we focused on the role and cellular target of the small phage protein Cg1978.

### Overproduction of Cg1978 triggers the activation of SOS response and RecA-independent prophage induction

As the CGP3 prophage was already characterized to be inducible in an SOS-dependent as well as in an SOS-independent manner (Helfrich et al., 2015; Nanda et al., 2014; Pfeifer et al., 2016), we determined the SOS-dependency of Cg1978-mediated prophage induction. To this end, the fluorescent outputs of different reporter strains were measured via flow cytometry in a time-resolved manner during cg1978 overexpression. Besides the wildtypic prophage reporter strain ATCC 13032::P*_lys_*-*eyfp*, a wildtypic SOS reporter strain ATCC 13032::P*recA*-*venus* as well as an SOS-deficient prophage reporter strain ATCC 13032 Δ*recA*::P*_lys_*-*eyfp* lacking the co-protease RecA – required for the induction of the host SOS response (Janion, 2008) – were used.

As described above, overexpression of cg1978 resulted in a similar growth phenotype of all reporter strains characterized by an elongated lag-phase (marked in grey) with subsequent wildtypic growth. The impaired cell growth under cg1978 overexpression conditions was confirmed by microscopy analysis of the prophage reporter strain. Increased levels of Cg1978 led to an elongated cell morphology and a small fraction of cells featuring a strongly increased output of the prophage reporter (Figure 2C, Video S1 and S2).

**Figure 2:**
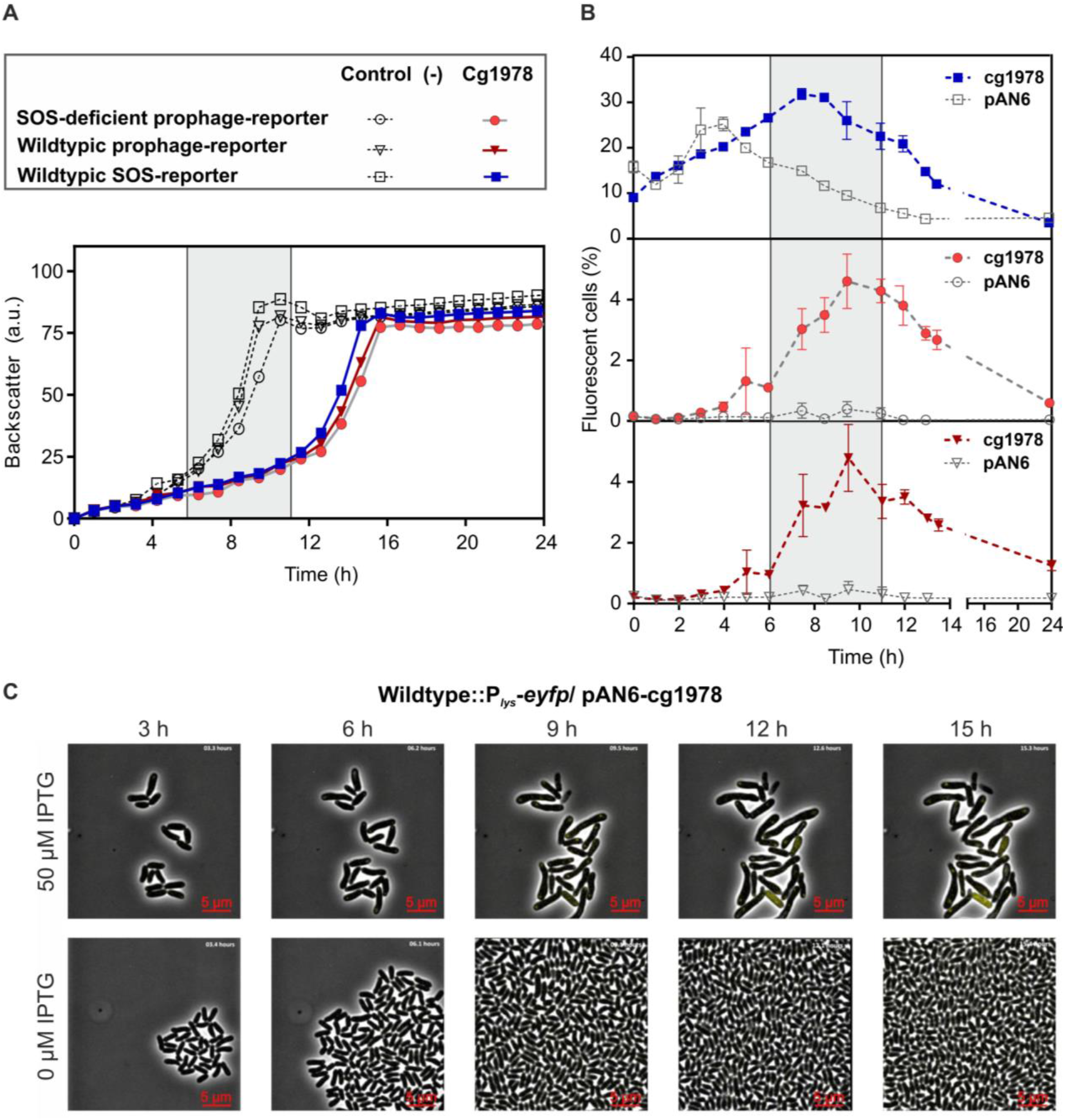
Time-resolved measurement of reporter outputs upon Cg1978 overproduction in *C. glutamicum* showed an activation of SOS response and RecA-independent prophage induction. Cultivation of a wildtypic SOS reporter strain ATCC 13032::P*_recA_*-*venus*, a wiltypic prophage reporter strain ATCC 13032::P*_lys_*-*eyfp* and an SOS-deficient prophage reporter strain ATCC 13032 Δ*recA*::P*_lys_*-*eyfp* carrying the plasmids pAN6 or pAN6-cg1978 was performed in the BioLector^®^ microcultivation system in CGXII-Kan_25_ medium with 2 % (w/v) glucose and 50 μM IPTG. All data represent mean values with standard deviations from three independent biological triplicates (n=3). **(A)** Growth curves based on the backscatter measurements in the BioLector^®^ microcultivation system. The elongated lag-phase of the Cg1978 overproducing strain is marked in grey. **(B)** Percentage of induced cells based on the flow cytometric measurements of eYFP or Venus fluorescence of the reporter strains. **(C)** Time-lapse fluorescence microscopy of the *C. glutamicum* prophage reporter strain ATCC 13032::P*_lys_*-*eyfp* carrying the pAN6-cg1978 plasmid. Cells were grown in PDMS-based microfluidic chip devices using CGXII medium supplemented with 25 μg ml^−1^ kanamycin. The medium was continuously supplied with a flow rate of 300 nl min^−1^ (Grünberger et al., 2015). Overexpression of cg1978 was induced by addition of 50 μM IPTG (Video S1 and S2).

Measurement of the reporter output over time confirmed CGP3 induction, but also revealed an induction of the cellular SOS response (Figure 2B). Interestingly, the wildtypic and the RecA-deficient prophage reporter strain showed nearly the same percentage of induced cells upon cg1978 overexpression throughout the entire measurement, reaching a peak value after 9.5 h of cultivation (Δ*recA*::P*_lys_*-*eyfp*: 4.6 ± 0.7, P*_lys_*-*eyfp*: 4.8 ± 0.9). These results emphasize CGP3 induction as a RecA-independent, indirect effect of cg1978 overexpression.

Remarkably, all reporter strains revealed an increasing fluorescence output during the lag-phase, which decreased again upon transition into the exponential growth phase (Figure 2B) suggesting the development of a subpopulation resistant to Cg1978 overproduction effects.

### Cg1978 directly interacts with gyrase subunit A (GyrA)

To identify the direct cellular target of Cg1978, we performed an in vitro pull-down assay. For this purpose, the small protein Cg1978 containing a C-terminal Strep-tag was overproduced in *E. coli* BL21 (DE3) and purified via affinity purification. The purified target protein was incubated with *C. glutamicum* cell extract and this sample was again passed over a Strep-Tactin column to identify proteins co-purifying with Cg1978. SDS PAGE analysis of proteins co-eluting with Cg1978 revealed an additional protein band at ~100 kDa (Figure 3A). Analysis of the co-purified protein via MALDI-TOF as well as analysis of the whole elution fraction via LC-MS/MS indicated the co-purification of Cg1978 with *C. glutamicum* (*C.g.*) DNA gyrase subunit A (~95 kDa), a subunit of the heterotetrameric ATP-dependent DNA gyrase complex (A_2_B_2_). This enzyme belongs to the subclass of Type II topoisomerases and plays a key role in DNA metabolism as it is able to introduce negative supercoiling to double-stranded DNA in an ATP-dependent manner (Reece & Maxwell, 1991).

**Figure 3:**
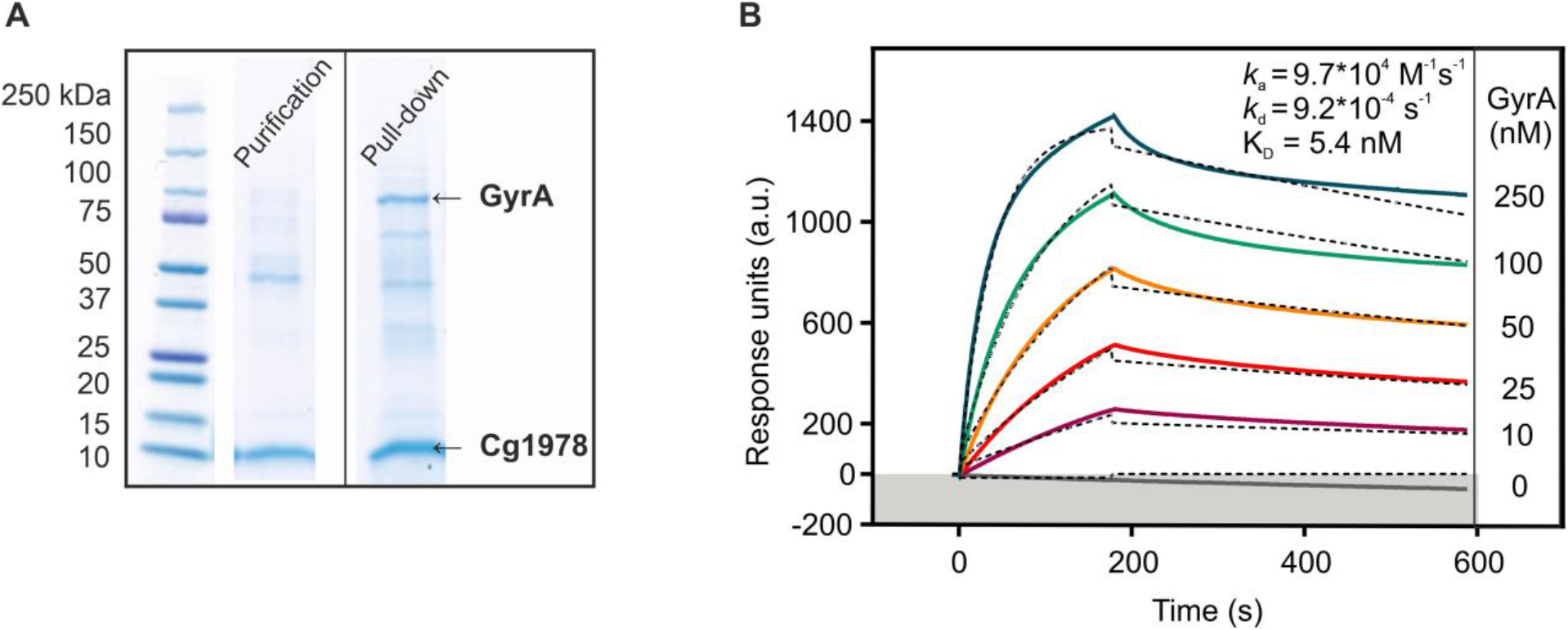
Cg1978 directly interacts with the gyrase subunit A (GyrA) in vitro. **(A)** The small protein Cg1978 containing a C-terminal Strep-tag was overproduced in *E. coli* BL21 (DE3) and purified via affinity purification. For the pull-down assay the *C. glutamicum* wildtype strain was cultivated in CGXII-Medium with 2 % (w/v) glucose until OD_600_ of 6. The purified target protein was incubated with *C. glutamicum* cell extract and again passed over a Strep-Tactin column aiming at the co-purification of Cg1978 with possible interaction partners. Proteins in the elution fractions were analyzed via SDS-PAGE and further identified using LC-MS/MS and MALDI-TOF. **(B)** Surface plasmon resonance spectroscopy of GyrA binding to Cg1978 (*k*_a_, association constant; *k*_d_, dissociation constant, K_D_, equilibrium dissociation constant). The colored lines represent the experimental data, the dotted lines represent the fitted data using 1:1 binding algorithm that were basis for the binding kinetics calculation.

As a next step, surface plasmon resonance spectroscopy was used to quantify the interaction between Cg1978 and GyrA by examining binding affinity and kinetics. The sensorgram revealed a stable and specific 1:1 interaction between Cg1978 and GyrA with a high association rate (*k*_a_=9.7×10^4^ M^−1^ s^−1^) and a slow dissociation rate (*k*_d_=9.2 x 10^−4^ s^−1^) resulting in an overall affinity (K_D_) of 5.4 nM (Figure 3B). Purification of Cg1978-C-His and GyrA-N-Strep for SPR analysis are shown in Supplementary Figure S3.

### Cg1978 inhibits DNA supercoiling via interaction with DNA gyrase in vitro

Due to its essential role for cell survival, DNA gyrase represents an important drug target of antibiotics and protein-based inhibitors (Collin et al., 2011). Based on the observed growth defect upon Cg1978 overexpression and the interaction with GyrA, we set out to further assess the effect of Cg1978 on gyrase activity by performing in vitro supercoiling inhibition assays with the purified enzyme.

For this purpose, Cg1978-C-Strep, GyrA-N-Strep and GyrB-C-Strep from *C. glutamicum* were purified separately using affinity chromatography (Figure 4A). Formation of the heterotetrameric enzyme complex was obtained by incubating equimolar amounts of both gyrase subunits on ice for 30 min. In a first step, the activity of the purified DNA gyrase was measured by incubating 0.5 μg relaxed pBR322 plasmid DNA with different gyrase concentrations. Addition of increasing gyrase concentrations led to an increasing supercoiling of the plasmid DNA resulting in a complete supercoiling using 200 nM of *C.g*. gyrase. This concentration was defined as 1 U and was used for testing the potential inhibitory effect of Cg1978 (Figure 4B).

**Figure 4:**
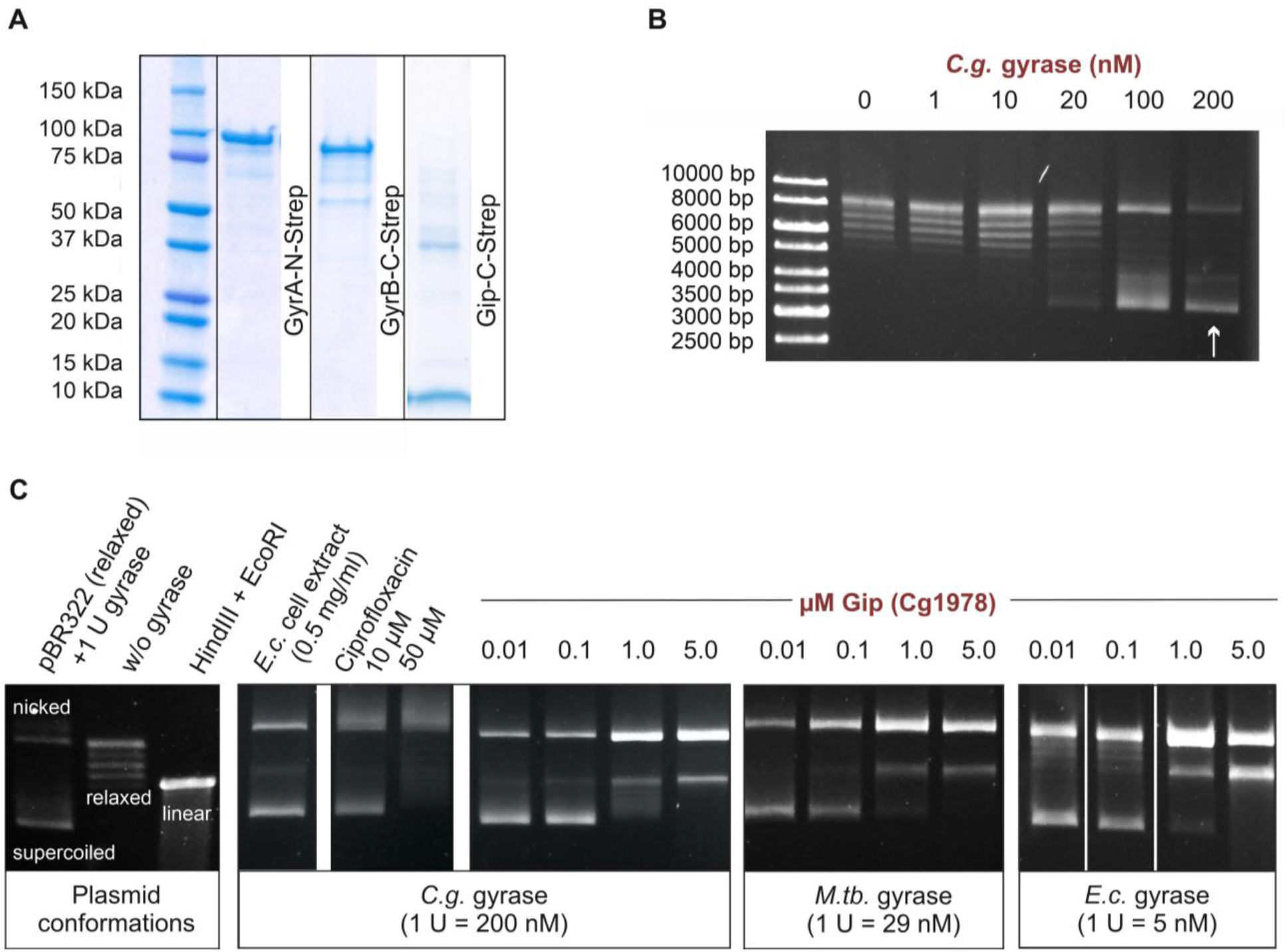
Cg1978 inhibits gyrase supercoiling activity in vitro. The assay was conducted according to the manual of the Gyrase Supercoiling Inhibition Assay-Kit from Inspiralis (Norwich, UK). **(A)** Gyrase subunit A containing a N-terminal and subunit B containing a C-terminal Strep-tag were separately overproduced in *E. coli* BL21 (DE3) and purified via affinity purification. The following gel electrophoresis was performed with 4 −20 % gradient gels at 120 V for 60 min. **(B)** After formation of the heterotetrameric gyrase complex, the activity assay of the purified DNA gyrase from *C. glutamicum* ATCC 13032 was performed to identify the gyrase concentration required for maximal supercoiling of 0.5 μg relaxed plasmid DNA, which was defined as 1 U. **(C)** Supercoiling inhibition assay to test the inhibitory effect of Cg1978 on 1 U of the DNA gyrases from *C. glutamicum* ATCC 13032, *E. coli* and *M. tuberculosis*. The *E. coli* BL21 (DE3) cell extract was further investigated regarding its inhibitory effect as this strain was used for protein overproduction. The known gyrase inhibitor Ciprofloxacin served as a positive control for efficient gyrase inhibition.

As shown in Figure 4C, incubation of increasing concentrations of Cg1978 (0.01-5.0 μM) with 1 U DNA gyrase and relaxed plasmid DNA resulted in a decrease in supercoiling activity of the *C.g*. DNA gyrase. A complete inhibition of the overall supercoiling activity was achieved by addition of 5 μM Cg1978. Therefore, we termed the gene product of cg1978 as Gip (gyrase inhibiting protein). To further investigate the activity profile of Gip, we determined its inhibitory activity on the DNA gyrases of *Mycobacterium tuberculosis* and *E. coli*. Gyrase subunits A of the actinobacterial strains *C. glutamicum* and *M. tuberculosis (M.tb.*) share a sequence identity of ~71%, while GyrA of *C. glutamicum* and the enterobacterial strain *E. coli (E.c.*) only show ~45 % sequence identity (Supplementary Figure S4and S5).

As described previously for the DNA gyrase of *C. glutamicum*, 1 U of the *E.c*. and *M.tb*. gyrases were used to examine supercoiling inhibition via Gip. The supercoiling assay revealed comparable inhibitory concentrations of Gip for the *E.c*. and *M.tb*. gyrase as for the DNA gyrase from *C. glutamicum*. Accordingly, for all tested gyrases 5 μM Cg1978 were successful to achieve a complete inhibition of DNA supercoiling, which results in an accumulation of the linear as well as the relaxed/nicked plasmid conformation (Figure 4C).

The inhibition of the *E.c*. gyrase by Gip was further in line with the fact that overexpression of gip resulted in a marked growth defect of *E. coli* in comparison to the empty vector control (Supplementary Figure S6).

To exclude, that the inhibitory effect of Gip on the DNA gyrase is caused by any other protein unspecifically co-purified from *E. coli* BL21 (DE3), the total cell extract of this strain was tested regarding its inhibitory effect. No influence on gyrase activity could be observed upon addition of the *E. coli* cell extract. As a positive control, the known fluoroquinolone-based gyrase-inhibitor ciprofloxacin was used and showed a complete inhibition of the supercoiling activity of the *C.g*. gyrase at 50 μM (16.6 μg ml^−1^). This was in line with already published data for the DNA gyrase of *M. smegmatis*, which showed 50 % inhibition of the supercoiling activity of 1 U gyrase by addition of 10 μg ml^−1^ ciprofloxacin (Manjunatha et al., 2002). In contrast to Gip where we also observed the accumulation of linear plasmid DNA, addition of ciprofloxacin just showed relaxed/nicked plasmid conformation in this assay.

### *Compensatory expression of* gyrAB *and* topA *as response to gyrase inhibition via Gip*

Supercoiling inhibition assays showed that Cg1978 (Gip) is a gyrase-inhibiting protein. As inhibition of the gyrase is lethal for the cell, we wanted to investigate how *gip* overexpression influences genome-wide expression levels. Since the overexpressing strain *gip* revealed a wildtypic growth rate after an elongated lag-phase, we were especially interested to understand how the bacterial host did counteract gyrase inhibition caused by Gip overproduction. To this means, comparative transcriptome analysis of the *C. glutamicum* ATCC 13032 strain containing the empty vector control and the strain containing the overexpression plasmid pAN6-*gip* was performed using DNA microarrays. The shown transcriptomic changes are based on mRNA levels of cells harvested at the same OD_600_ of 6 in the mid-exponential growth phase.

The *gip* overexpressing strain showed partially upregulation of CGP3 genes due to overexpression of *gip* (Supplementary Table S1), confirming the prophage induction also revealed by the above described reporter assays. Apart from the CGP3 region, overexpression of *gip* led to upregulation of 352 genes and downregulation of 334 genes reflecting the high impact of gyrase inhibition on overall cell metabolism.

Interestingly, both gyrase subunits *gyrA* and *gyrB* were markedly upregulated showing more than 4-fold increase in expression levels (Table 1). In contrast, *topA* coding for topoisomerase I, which catalyzes the opposing reaction of the DNA gyrase by removing negative supercoils, showed a reduced expression level. Also expression of further genes involved in DNA metabolism was influenced by *gip* overexpression including exemplarily the downregulation of different genes coding for helicases (cg0838, cg0842, cg0843, cg0845, cg0889 and cg1498). Additionally, nine targets of the SOS key player LexA, e.g. *recN* (DNA repair) and *ftsK* (cell division and chromosome segregation), showed an increased mRNA ratio, which was in line with the high fluorescent outputs of the SOS reporter strain upon Gip overproduction (Table 1). A complete overview on the transcriptomic changes is shown in Supplementary Table S4.

**Table 1:** Impact of *gip* overexpression on global expression levels. A genome-wide comparison of mRNA levels the *C. glutamicum* ATCC 13032 strain overexpressing *gip* and the wildtype strain carrying the empty vector control was performed. The shown mRNA ratios indicate mean values from three independent biological replicates (n=3). The strains were cultivated in CGXII medium with 2 % (w/v) glucose and 50 μM IPTG and mRNA was prepared from cells at an OD_600_ of 6. The mRNA ratios were calculated by dividing the mRNA levels of *gip* overexpressing strain by the mRNA levels of the strain carrying the empty vector control. The table includes selected genes which showed a changed mRNA level in all experiments (mRNA ratio > 2.0: upregulation (red) or < 0.5: downregulation (green), p-value ≤ 0.05).

| Gene locus | Gene name | Annotation | mRNA ratio<sup>a</sup> |
| --- | --- | --- | --- |
| **Genes coding for proteins involved in DNA metabolism** |  |  |  |
| cg0015 | *gyrA* | DNA gyrase subunit A, DNA topoisomerase II | 6.23 |
| cg0007 | *gyrB* | DNA gyrase subunit B, DNA topoisomerase II | 4.43 |
| cg0373 | *topA* | DNA topoisomerase I | 0.40 |
| cg0845 |  | superfamily II DNA/RNA helicase, SNF2 family | 0.49 |
| cg0889 |  | putative DNA helicase RecQ | 0.48 |
| cg0843 |  | helicase | 0.45 |
| cg1498 |  | RecG-like helicase | 0.43 |
| cg0842 |  | putative DNA helicase | 0.39 |
| cg0838 |  | helicase | 0.22 |
| **LexA target genes** |  |  |  |
| cg1602 | *recN* | DNA repair protein | 10.96 |
| cg1255 |  | putative HNH endonuclease, conserved | 5.11 |
| cg0470 | *htaB* | secreted heme transport-associated protein | 4.41 |
| cg0738 | *dnaE2* | DNA polymerase III subunit $\alpha$ (EC:2.7.7.7) | 3.11 |
| cg1288 |  | putative multidrug efflux permease of the major facilitator superfamily | 2.83 |
| cg1080 |  | putative multicopper oxidase | 2.67 |
| cg2158 | *ftsK* | cell division protein | 2.46 |
| cg0713 |  | hypothetical protein, conserved | 2.13 |
| cg2114 | *lexA* | transcriptional regulator, involved in SOS/stress response, LexA-family | 2.09 |
| cg2950 | *radA* | DNA replication, recombination, repair, and degradation | 0.48 |
| cg2381 |  | Unknown function | 0.47 |
| cg0834 | *tusE* | Carbon source transport and metabolism | 0.34 |
| cg0841 |  | Unknown function | 0.31 |
| cg1314 | *putP* | Amino acid transport and metabolism | 0.30 |
| cg3345 |  | Unknown function | 0.24 |

<sup>a</sup> The mRNA ratios were calculated by dividing the mRNA levels of *gip* overexpressing strain by the mRNA levels of the strain carrying the empty vector control.

## Discussion

In this study, a screening of small cytoplasmic proteins encoded by the CGP3 prophage of *C. glutamicum* resulted in the identification of Gip (Cg1978, 6.8 kDa) representing a novel gyrase inhibitor protein. Overproduction of Gip resulted in significant growth defects and prophage induction in a subpopulation. Further characterization of this small phagic protein confirmed a specific, stable and high affinity interaction with the GyrA subunit and an inhibition of the supercoiling activity of the DNA gyrase in vitro.

The DNA gyrase possesses the unique ability to catalyze the ATP-dependent negative supercoiling of double-stranded DNA by cleaving and rejoining (Bush et al., 2015). Supercoiling inhibition assays showed that Gip-mediated gyrase inhibition resulted in accumulation of the linear plasmid conformation, while no more supercoiled plasmid DNA was detectable (Figure 4). This observation suggests a similar mode of action like synthetic fluoroquinolones and the well-characterized proteinaceous bacterial toxins Microcin B17 (MccB17, 3.1 kDa) and CcdB (11.7 kDa), which poison the DNA gyrase by stabilizing the gyrase-DNA cleavage complex leading to double strand DNA breaks (Bernard et al., 1993; Drlica & Malik, 2003; Pierrat & Maxwell, 2003). In contrast, natural-derived aminocoumarins (e.g. novobiocin) inhibit ATP hydrolysis as the binding site overlaps with the ATP binding pocket of the GyrB subunit (Maxwell & Lawson, 2003). Remarkably, the tested gyrases of *E. coli, M. tuberculosis* and *C. glutamicum* showed a very similar susceptibility to Gip-mediated inhibition. In all cases a complete inhibition of supercoiling was achieved upon addition of 5 μM Gip in in vitro assays. In contrast, the inhibitory activity of CcdB and MccB17 were described to be specific to the DNA gyrase of their bacterial host *E. coli* as the gyrase of *M. smegmatis* was not poisoned by these proteins in vitro (Chatterji et al., 2001). Even though the DNA gyrase is a conserved protein among bacteria, Gram-positive and Gram-negative bacteria show substantial differences in the amino acid sequence of GyrAB (Madhusudan & Nagaraja, 1996; Manjunatha et al., 2000). Accordingly, the absence of specific target residues potentially explains the different levels of susceptibility of the *E. coli* and *M. smegmatis* gyrase to the CcdB and MccB17 toxins (Chatterji et al., 2001). In contrast to this, supercoiling inhibition assays performed in this study suggested that Gip targets a conserved fold in the GyrA subunit. However, further studies and structural analysis are required to elucidate the exact molecular mechanism of Gip-mediated gyrase inhibition.

In line with the proposed mode of action, reporter outputs of the RecA-dependent SOS reporter strain and transcriptomic analysis of Gip overproducing cells revealed an induction of the SOS response (Figure 2B, Table 1). This finding conforms to already published data describing an activation of SOS response as one of the pleiotropic effects of gyrase inhibition (Jeong et al., 2006). Stabilization of the gyrase-cleaved DNA complex results in arrested replication forks and widespread DNA damage by stimulating the formation of DNA breaks triggering the SOS response (DeMarini & Lawrence, 1992; Dwyer et al., 2007).

Gip overproduction was further shown to activate the induction of the CGP3 prophage. However, the fact that the observed growth defect of Gip overproducing cells is independent of the presence of the CGP3 prophage (Figure 1B) and that deletion of *gip* did not result in an altered inducibility of CGP3 (Figure S2) emphasized prophage activation as an indirect effect of Gip overproduction. Recent studies already confirmed that CGP3 is inducible in an SOS-dependent as well as in an SOS-independent manner (Helfrich et al., 2015; Nanda et al., 2014; Pfeifer et al., 2016). As the wildtypic and the RecA-deficient prophage reporter strain revealed nearly identical fluorescent outputs, we propose that prophage induction occurred mainly in a RecA-independent manner. Here, changes in DNA topology due to gyrase inhibition might be a possible reason for CGP3 induction. The lysogenic state of CGP3 is maintained by the Lsr2-type silencer protein CgpS, which was shown to target more than 35 AT-rich regions within the CGP3 element (Pfeifer et al., 2016). The formation of this dense nucleoprotein complex was shown to be crucial for efficient CgpS-mediated silencing (Wiechert et al., 2020). Especially in case of proteins targeting AT-rich DNA sequences, the topological state of DNA can affect protein-DNA interactions (Dorman & Dorman, 2016). Apart from that, different studies already demonstrated an influence of DNA topology on the λ repressor and the lysogenic-lytic decision of phage λ (Ding et al., 2014; Norregaard et al., 2013, 2014).

Regarding the native function of Gip, we hypothesize that encoding a gyrase inhibitor could be advantageous for the phage as it might enable a more efficient phage DNA replication by modulating gyrase activity. Similar assumptions were previously reported for the protein 55.2 encoded by the T4 phage of *E. coli*. By modulating DNA relaxation activity of TopoI, gp55.2 was proposed to be required for an optimal phage yield during infection (Mattenberger et al., 2015).

Global topological alterations caused by Gip overproduction were also reflected by the transcriptome analysis revealing a marked impact on global gene expression patterns (Table 1). Particularly noteworthy in this context are the significantly increased mRNA levels of *gyrA* and *gyrB* as well as the downregulation of *topA*. Since the DNA topology modulatory proteins, gyrase and topoisomerase I (TopoI), catalyze opposing reactions of DNA supercoiling and relaxation, the observed effect likely reflects compensatory mechanisms ensuring DNA topology homeostasis (Massé & Drolet, 1999; Reece & Maxwell, 1991). As the DNA gyrase is indispensable for replication and transcription by changing the topological state of DNA, its inhibition was previously described to globally affect the gene expression profile (Guha et al., 2018; Jeong et al., 2006). Previous studies showed further that expression of the *gyrAB* and *topA* is controlled in a supercoiling-sensitive manner: While increasing DNA relaxation stimulates *gyrAB* expression (Menzel & Gellert, 1983), it represses expression of *topA* allowing a homeostatic maintenance of DNA topology (Ahmed et al., 2016). As gyrase inhibition causes DNA relaxation, increased expression levels of *gyrAB* and decreased expression level of *topA* are a well-known compensatory mechanism of gyrase inhibition (Cheung et al., 2003; Jeong et al., 2006; Menzel & Gellert, 1983). The adaptation at the level of gene expression could then explain the resumed growth of the *gip* expressing strain – reaching almost wildtypic growth rates after a pronounced lag phase (Figure 1).

In summary, we have identified Gip as a novel gyrase inhibitor protein encoded by the CGP3 prophage of *C. glutamicum*. In vitro experiments provided evidence for a similar mode of action of Gip compared to the bacterial toxin CcdB and microcin B17 by stabilizing the gyrase-DNA cleavage complex via interaction with GyrA – thereby preventing DNA supercoiling. Activity profiling of Gip revealed a broad susceptibility of different gyrases from Gram-positive and Gram-negative bacteria. The mycobacterial DNA gyrase has already been used as a target for anti-mycobacterial therapy and fluoroquinolones show only limited success as second-line drugs due to increasing multidrug-resistance (Falzon et al., 2013). Therefore, a detailed elucidation of the mechanism of action of Gip may permit the design of gyrase inhibitors with a broad activity profile.

## Material and methods

### Bacterial strains and growth conditions

All bacterial strains and plasmids used in this study are listed in Supplementary Table S1 and S2, respectively. *Corynebacterium glutamicum* ATCC 13032 (NCBI reference: NC_003450.3) was used as wild type strain (Ikeda & Nakagawa, 2003). For growth studies and fluorescence measurements as well as for transcriptome analysis, *C. glutamicum* cells were pre-cultivated in BHI (brain heart infusion, Difco BHI, BD, Heidelberg, Germany) at 30 °C and 170 rpm for 8 hours. The pre-culture was used to inoculate an overnight-culture in CGXII minimal medium with 2 % (w/v) glucose, which was cultivated under the same conditions. The next day, the overnight-culture was finally used to inoculate the main culture in CGXII minimal medium with 2 % (w/v) glucose to an OD_600_ of 1. To all media, kanamycin was added in a concentration of 25 μg ml^−1^. Gene expression was induced using 50 μM IPTG (Isopropyl β-D-1-thiogalactopyranoside). For standard cloning applications, *E. coli* DH5α was cultivated in Lysogeny Broth (Difco LB, BD, Heidelberg, Germany) medium containing 50 μg ml^−1^ kanamycin (LB Kan_50_) at 37°C and 120 rpm. For protein overproduction and following purification, *E. coli* BL21 (DE3) strain was used. Pre-cultivation was performed in LB Kan_50_ medium, which was incubated overnight at 37 °C and 120 rpm. The main culture in LB Kan_50_ medium was inoculated to an OD_600_ of 0.1 using the pre-culture. At an OD_600_ of 0.6 gene expression was induced using 100 μM IPTG. Cells were harvested after additional 24 h incubation at 16 °C.

### Recombinant DNA work and cloning techniques

All plasmids and oligonucleotides used in this study are listed in Supplementary Table S2 and S3, respectively. Standard cloning techniques like PCR and restriction digestion were performed according to standard protocols (Sambrook & Russell, 2001). In all cases, Gibson assembly was used for plasmid construction (Gibson, 2011). DNA regions of interest were amplified via PCR using the chromosomal DNA of *C. glutamicum* ATCC 13032 as template. The plasmid backbone was cut using the indicated restriction enzymes. Sequencing and synthesis of oligonucleotides was performed by Eurofins Genomics (Ebersberg, Germany). Genomic deletions were constructed using the pK19*mobsacB* plasmid and the two-step homologues recombination method (Niebisch & Bott, 2001). The 500 bp up- and downstream regions of the respective gene were amplified using the oligonucleotides listed in Supplementary Table S3. Both PCR products and the digested pK19*mobsacB* plasmid (with BamHI, EcoRI) were assembled via Gibson assembly (Gibson, 2011). Correct deletion was verified by sequencing of the colony PCR product with the indicated oligonucleotides (Supplementary Table S3).

### Microtiter cultivation and reporter assays

For growth experiments and fluorescence assays, the BioLector^®^ microcultivation system of m2p-labs (Aachen, Germany) was used (Kensy et al., 2009). The main cultivation was executed in FlowerPlates (MTP-48-B, m2p-labs) at 30 °C and 1200 rpm as described in (Heyer et al., 2012) using 750 μl of CGXII-Kan_25_ minimal media with 2% (w/v) glucose containing 50 μM IPTG and 25 μg ml^−1^ kanamycin. During cultivation, biomass was measured as a function of backscattered light intensity with an excitation wavelength of 620 nm (filter module: λ_Ex_/ λ_Em_: 620 nm/ 620 nm, gain: 15), while fluorescence intensity was recorded with an excitation wavelength of 488 nm and an emission wavelength of 520 nm (filter module: λ_Ex_/ λ_Em_: 488 nm/ 520 nm, gain: 60). The measurements of backscatter and fluorescence were taken in 15 min intervals.

### Protein purification via affinity tags

For heterologous protein overproduction, *E. coli* BL21 (DE3) cells containing the pET-cg1978-C-*strep* plasmid, the pET-*gyrA*-N-*strep* plasmid, the pET-*gyrB*-C-*strep* plasmid or the pET-cg1978-*C*-*his* plasmid were cultivated as described in bacterial strains and growth conditions.

Cell harvesting and disruption were performed as described by (Pfeifer et al., 2016) using buffer A (100 mM Tris-HCl, pH 8.0) for cell disruption and buffer B (100 mM Tris-HCl, 250 mM NaCl, pH 8.0) for purification. Purification of Strep-tagged Cg1978 was conducted by applying the supernatant to an equilibrated 2 ml Strep-Tactin^®^-Sepharose^®^ column (IBA, Göttingen, Germany). After washing with 20 ml buffer B, the protein was eluted with 6 ml buffer B containing 15 mM d-desthiobiotin (Sigma Aldrich). Purification of GyrA-N-Strep and GyrB-C-Strep were conducted in the same way using an adjusted buffer B*_gyr_* for cell disruption and purification (buffer B*_gyr_*: 20 mM Tris-HCl, 500 mM NaCl, 10 % (w/v) glycerol, 1 mM DTT, pH 7.9).

For purification of Cg1978-C-His, the cell pellet was resuspended in 50 ml TNI buffer (20 mM Tris-HCl, 20 mM imidazole, pH 8.0) and cells were disrupted as described above. Purification of His-tagged Cg1978 was performed by applying the supernatant to an equilibrated 2 ml Ni-NTA column (Invitrogen, Kalifornien, USA). After washing with 30 ml TNI20 buffer, the protein was eluted with increasing imidazole concentrations using TNI buffer (20 mM Tris-HCl, pH 8.0) containing 50 mM, 100 mM, 200 mM or 400 mM imidazole.

After every purification, the elution fractions with the highest protein concentration were pooled and analyzed with SDS-PAGE (Laemmli, 1970) using a 4–20% Mini-PROTEAN^®^ gradient gel (Bio Rad, Munich, Germany).

### In vitro pull-down assay and MALDI-TOF analysis

Protein purification of Cg1978-C-Strep was conducted as described above. The elution fractions showing the highest protein concentration in a Bradford-assay (Bradford, 1976) were pooled and purified with size-exclusion chromatography using PD10 desalting columns (GE Healthcare) and buffer B (100 mM Tris-HCl, 250 mM NaCl, pH 8.0) according to manufacturer’s manual to remove excess desthiobiotin. For the detection of possible interaction partners of the target protein on a protein-protein level*, C. glutamicum* ATCC 13032 wildtype cells were cultivated in BHI medium. At an OD_600_ of 5-6 the cells were harvested at 11.325 *g* and 4 °C for 15 min and cell pellet of 100 ml cell culture was resuspended in 25 ml buffer A (100 mM Tris-HCl, pH 8.0) with cOmplete™ Proteinase inhibitor (Roche). Cell disruption was performed using the French Press cell with a pressure of 172 mPA for five passages, followed by a centrifugation step at 5000 *g* for 50 min.

For co-purification of possible protein interaction partners, the purified target protein Cg1978 was incubated with the *C. glutamicum* crude extract at RT for 1 h. After loading the mixture to the StrepTactin column, the purification was performed as described above.

The elution fractions with the highest protein concentration were precipitated by addition of 100 % (w/v) trichloroacetic acid (TCA) in a volume ratio of 4 units protein to 1 unit TCA (Sivaraman et al., 1997). After incubation at 4 °C overnight the precipitation approach was centrifuged for 5 min at 14.000 g. The supernatant was discarded and the pellet was washed with 200 μl cold acetone twice. Afterwards the pellet was dried for 10 min at 95 °C and resuspended in 30 μl 1.5 x SDS loading buffer for gel electrophoresis or in 30 μl trypsin reaction buffer provided by the Trypsin Singles, Proteomics Grade kit (Sigma-Aldrich) for LC-MS/MS sample preparation.

Analysis of elution fractions via SDS-PAGE (Laemmli, 1970) was performed using a 4–20% Mini-PROTEAN^®^ gradient gel (Bio Rad, Munich, Germany). After staining of the gels with Coomassie dye based RAPIDstain solution (G-Biosciences, St. Louis, MO, USA) MALDI-TOF-MS measurements were performed with an Ultraflex III TOF/TOF mass spectrometer (Bruker Daltonics, Bremen, Germany) for identification of the co-purified proteins (Bussmann et al., 2010). Elution fractions were further analyzed via LC-MS/MS.

### LC-MS/MS sample preparation and measurement

LC-MS/MS after TCA precipitation was executed using the Trypsin Singles, Proteomics Grade kit (Sigma-Aldrich) according to the manufacturer’s instruction. The prepared tryptic peptide samples were separated chromatographically on a nanoLC Eksigent ekspert™ 425 LC system in microLC modus (Sciex) coupled with a 25 Micron ESI Electrode to a TripleTof™ 6600 mass spectrometer (Sciex). As a trap, a YMC-Triart C18 column with the dimension 5 x 0.5 mm ID, 3 μm, 12 nm (YMC) was used, combined with a YMC-Triart C18 column with 150 x 0.3 mm ID, 12 nm, S-3μm (YMC) as analytical column. The column oven was set to 40°C.

For trapping, 2% acetonitrile in dd.H_2_O with 0.5% formic acid served as a loading solvent, while 0.1 % formic acid was used as mobile phase A and acetonitrile with 0.1% formic acid (both LC-MS grade, ROTISOLV^®^, ≥ 99.9%, Carl Roth) as mobile phase B. First, 10 μl of each sample containing up to 8 μg of digested protein was loaded from the cooled autosampler onto the trap column using 100% loading solvent for 10 minutes at 10 μl min^−1^ for desalting and enrichment.

For the following separation of the peptides on the analytical column a linear gradient with increasing concentrations of mobile phase B was used starting with 97 % A and 3 % B and a flow rate of 5 μL min^−1^ as initial condition. During Information Dependent Acquisition (IDA) and SWATH measurements the following source and gas settings were applied: 5500 V spray voltage, 35 psi curtain gas, 12 psi ion source gas 1, 20 psi ion source gas 2, and 150°C interface heater. Each sample was injected three times.

For shotgun measurements the mass spectrometer was operated with a “top 50” method: Initially, a 250 ms survey scan (TOF-MS mass range m/z 400-1500, high-resolution mode) was collected from which the top 50 precursor ions were automatically selected for fragmentation, whereby each MS/MS 97 Appendix event (mass range m/z 170-1500, in high sensitivity mode) consisted of a 40 ms fragment ion scan. For parent ion selection, the precursor ion intensity served as the main selection criterion. Ions with an intensity exceeding 100 counts per second and with a charge state of 2+ to 5+ were preferentially selected. Selected precursors were added to a dynamic exclusion list for 22 s and subsequently isolated using a quadrupole resolution of 0.7 amu and fragmented in the collision cell with a rolling collision energy (CE) of 10 eV. If less than 50 precursor ions fulfilling the selection criteria were detected per survey scan, the detected precursors were subjected to extended MS/MS accumulation time to maintain a constant total cycle time of 2.3 s.

For data analysis the IDA data were processed with ProteinPilot™ (V5.01, Sciex, USA) using the ParagonTM Algorithm for peptide identification and the ProGroup™ Algorithm for protein identification.

### DNA microarrays

For a comparative transcriptome analysis of *C. glutamicum* ATCC 13032 carrying the empty pAN6 vector and cells carrying the pAN6-cg1978 vector, cultivation was performed as described in ‘Bacterial strains and growth conditions’ using CGXII minimal media with 2% (w/v) glucose and 50 μM IPTG. For both strains, cells were harvested at an OD_600_ of 6. RNA purification was done using the “RNeasy Mini”-Kit (QIAGEN) according to the manufacturer’s manual. The preparation of labeled cDNA and DNA microarray analysis was performed as described previously (Donovan et al., 2015). The data processing was executed with an in-house software according to (Polen & Wendisch, 2004). Genes with an mRNA ratio (sample/neg. control) > 2.0 (p-value < 0.05) were classified as upregulated, whereas genes with an mRNA ratio < 0.5 (p-value < 0.05) were classified as downregulated. Array data were deposited in the GEO database (ncbi.nlm.nih.gov/geo) with accession number GSE151224.

### Flow cytometry

Analysis of fluorescent reporter outputs at the single cell level was performed using the BD Accuri™ C6 Plus flow cytometer (BD Bioscience, Germany). The chromophore of the yellow fluorescent protein eYFP or Venus was excited with a blue dye laser with an excitation wavelength of 488 nm. Fluorescence emission of eYFP and Venus was measured using a 530/30 nm standard filter. Particle size was detected using the forward light scatter (FSC). The flow cytometer was started up by flushing with filtered, dd. H_2_O for 10 min. For preparing flow cytometry samples, cell cultures were mixed with 1 ml flow cytometric fluid (BD™ 342003 FACS-flow Sheath Fluid). The concentration of the samples was adjusted to a measurement of 2.000 to 8.000 events/s. Flow cytometric data were analyzed using FlowJo V10 software (Tree Star, Ashland, OR).

### Cultivation and perfusion in microfluidic device

Single-cell analysis of cg1978 overexpressing cells was performed using an in-house developed microfluidic platform (Grünberger et al., 2015; Grünberger et al., 2013; Helfrich et al., 2015). Cultivation and time-lapse imaging was performed in CGXII medium with 2% (w/v) glucose and 25 μg ml^−1^ kanamycin as described by (Pfeifer et al., 2016). Overexpression of cg1978 in the prophage reporter strain ATCC 13032::P*_lys_*-*eyfp* was induced by adding 50 μM IPTG to the medium. An uninduced control served as a reference.

### Supercoiling inhibition assay

For the supercoiling inhibition assay, Cg1978 as well as both gyrase subunits (GyrA and GyrB) were purified by the means of a C-terminal Strep-Tag for Cg1978 and GyrB and an N-terminal Strep-Tag for GyrA as described above. Using PD10 desalting columns (GE Healthcare, Illinois, U.S) the buffer of Cg1978 was exchange to PBS (137 mM NaCl, 2.7 mM KCl, 8 mM Na2HPO4, 1.5 mM KH2PO4, pH 7.4). In case of GyrA and GyrB, the buffer was exchanged to 50 mM Tris-HCl, pH 7.9, 30 % (w/v) glycerol, 5 mM DTT. Formation of the heterotetramic gyrase complex was obtained by incubating equimolar amounts of GyrA and GyrB for 30 min on ice. The activity of the purified *C. glutamicum* (*C.g.*) gyrase as well as its inhibition by Cg1978 were determined according to the manufacturer’s manual of the Gyrase (HIS) Supercoiling Assay Kits (Inspiralis, Norwich, UK). According to their assay conditions, the *C.g*. gyrase concentration of 200 nM resulting in complete supercoiling of 0.5 μg relaxed plasmid DNA was determined as 1 U. Supercoiling inhibition of *C.g*. gyrase was assayed by using 1 U of the *C.g*. gyrase with increasing concentrations of Cg1978 (0.01-5.0 μM). Additionally, different concentrations of the known inhibitor ciprofloxacin (10 and 50 μM) were used as positive control. The *E. coli* BL21 (DE3) cell extract was further tested as a negative control since this host was used for Cg1978 protein overproduction.

Moreover, the inhibitory effect of Cg1978 on the *M.tb* and *E.c*. gyrase were investigated according to the *M. tuberculosis and E. coli* Gyrase (HIS) Supercoiling Assay Kit (Inspiralis, Norwich, UK) using 1 U of the respective gyrases and the same Cg1978 concentrations as for the *C.g*. gyrase.

### Surface Plasmon Resonance Spectroscopy (SPR)

For SPR analysis, Cg1978-C-His and GyrA-N-Strep were purified as described above. After purification, the buffer of both proteins was exchanged to PBS (137 mM NaCl, 2.7 mM KCl, 8 mM Na2HPO4, 1.5 mM KH2PO4, pH 7.4) using PD10 Desalting columns (GE Healthcare, Illinois, U.S). Binding of His-tagged Cg1978-C-His to GyrA-N-Strep was analyzed by SPR analysis in a Biacore 3000 device (GE Healthcare, Freiburg, Germany) using a Sensor Chip CM5 (GE Healthcare, Freiburg, Germany). As first step, an anti-histidine antibody (GE Healthcare, Freiburg, Germany) was immobilized to the chip matrix using amino coupling chemistry. All experiments were carried out in HBS-EP buffer (0.01 M HEPES, pH 7.4, 0.15M NaCl, 3 mM EDTA, 0.005% v/v Surfactant P20) at 25°C. Following the standard coupling protocol for antibody immobilization the mixture of 0.05 M N-Hydroxysuccinimide (NHS) and 0.2 M 1-Ethyl-3-(3-dimethylaminopropyl) carbodiimide hydrochloride (EDC) was injected for a total contact time of 420 s to activate the matrix. Then, the anti-histidine antibody (50 μg ml^−1^) diluted in immobilization buffer (10 mM sodium acetate, pH 4.5) was injected for 420 s. To deactivate the unbound parts of the chip matrix, 1 M ethanolamine hydrochloride-NaOH (pH 8.5) was injected for 420 s. The flow rate was set to 10 μl min^−1^ during this immobilization procedure. Approximately 8.000-10.000 response units (RU) of the anti-histidine antibody were immobilized per flow cell. For the binding analysis, 180-250 RU of Cg1978-C-His was captured via injection of 40 μl (10 nM) at a flow rate of 5 μl min^−1^ followed by 10 min of HBS-EP buffer to remove unbound protein from the chip. The binding analysis between Cg1978-C-His and GyrA-N-Strep was then performed by injecting 90 μl of GyrA-N-Strep (10-250 nM) followed by a dissociation time of 300 s at a flow rate of 30 μl min^−1^. After each cycle, the surface was regenerated by injection of regeneration buffer (10 mM Glycine-HCl, pH 1.5) for 30 s, at a flow rate of 30 μl min^−1^. After the equilibration with three start up cycles without the analyte, this was repeated for various concentrations of GyrA-N-Strep (10 nM-250 nM). Sensorgrams were recorded using Biacore 3000 Control Software 4.1.2 and analyzed with BIAevaluation software 4.1.1 (GE Healthcare, Freiburg, Germany). The surface of flow cell 1 immobilized with the anti-histidine antibody was used to obtain blank sensorgrams for subtraction of the bulk refractive index background. The referenced sensorgrams were normalized to a baseline of 0. Peaks in the sensorgrams at the beginning and the end of the injection are due to the run-time difference between the flow cells for the chip.

## Acknowledgments

We thank the European Research Council (ERC Starting Grant, grant number 757563) and the Helmholtz Association (grant number W2/W3-096) for financial support.

We thank Astrid Wirtz for technical assistance with respect to the LC-MS/MS measurement.

SPR analyses were performed in the Bioanalytics service unit of the JGU Biocenter.

## Conflict of interest

The authors declare no conflict of interest.

## Author contributions

LK, MH and JF conceived the study; LK and JB performed the experiments; LK, MH, JB, RH and JF analyzed the data; LK and JF wrote the manuscript. All authors reviewed and edited the manuscript.

## Supplementary Information

**Table S1:** Bacterial strains used in this study.

**Table S2:** Plasmids used in this study.

**Table S3:** Oligonucleotides used in this study.

**Table S4:** Impact of cg1978 overexpression on global expression levels.

**Figure S1:** Screening of small phagic proteins regarding their impact on cellular growth and CGP3 induction in *C. glutamicum*.

**Figure S2:** Cell growth and prophage inducibility in the Δcg1914 or Δcg1978 mutant strain.

**Figure S3:** Purification of proteins for surface plasmon resonance spectroscopy of protein-protein interaction.

**Figure S4:** Sequence alignment of the DNA gyrase subunit from *C. glutamicum* (C), *M. tuberculosis* (M) and *E. coli* (E).

**Figure S5:** Sequence alignment of the DNA gyrase subunit B from *C. glutamicum* (C), *M. tuberculosis* (M) and *E. coli* (E).

**Figure S6**: Cell growth of *E. coli* BL21 (DE3) under *gip* overexpression conditions

**Video S1:** Time lapse video of a *C. glutamicum* microcolony of the prophage reporter strain under cg1978 overexpression (50 μM IPTG).

**Video S2:** Time lapse video of a *C. glutamicum* microcolony of the prophage reporter strain under standard conditions (0 μM IPTG).

## Notes

### Competing Interest Statement

The authors have declared no competing interest.

