## Supplemental Information for "Identification of Gip as a novel phage-encoded gyrase inhibitor protein featuring a broad activity profile"

Running title: Phage-encoded DNA gyrase inhibitor

Larissa Kever[1], Max Hünnefeld[1], Jannis Brehm[2], Ralf Heermann[2] and Julia Frunzke[1]*

[1]Institute of Bio- und Geosciences, IBG-1: Biotechnology, Forschungszentrum Jülich, 52425 Jülich, Germany

[2]Institut für Molekulare Physiologie, Biozentrum II, Mikrobiologie und Weinforschung, Johannes-Gutenberg-Universität Mainz, 55128 Mainz, Germany

*Corresponding authors:

**Content**

**Figure S1:** Screening of small phagic proteins regarding their impact on cellular growth and CGP3 induction in *C. glutamicum.*

**Figure S2:** Cell growth and prophage inducibility in the Δcg1914 or Δcg1978 mutant strain.

**Figure S3:** Purification of proteins for surface plasmon resonance spectroscopy of protein-protein interaction.

**Figure S4:** Sequence alignment of the DNA gyrase subunit A from *C. glutamicum, M. tuberculosis* and *E. coli.*

**Figure S5:** Sequence alignment of the DNA gyrase subunit B from *C. glutamicum, M. tuberculosis* and *E. coli.*

**Figure S6**: Cell growth of *E. coli* BL21 (DE3) under *gip* overexpression conditions

**Table S1:** Bacterial strains used in this study.

**Table S2:** Plasmids used in this study.

**Table S3:** Oligonucleotides used in this study.

**Table S4:** Impact of *gip* (cg1978) overexpression on global expression levels.

**Video S1:** Time lapse video of a *C. glutamicum* microcolony of the prophage reporter strain under cg1978 overexpression (50 µM IPTG).

**Video S2:** Time lapse video of a *C. glutamicum* microcolony of the prophage reporter strain under standard conditions (0 µM IPTG).

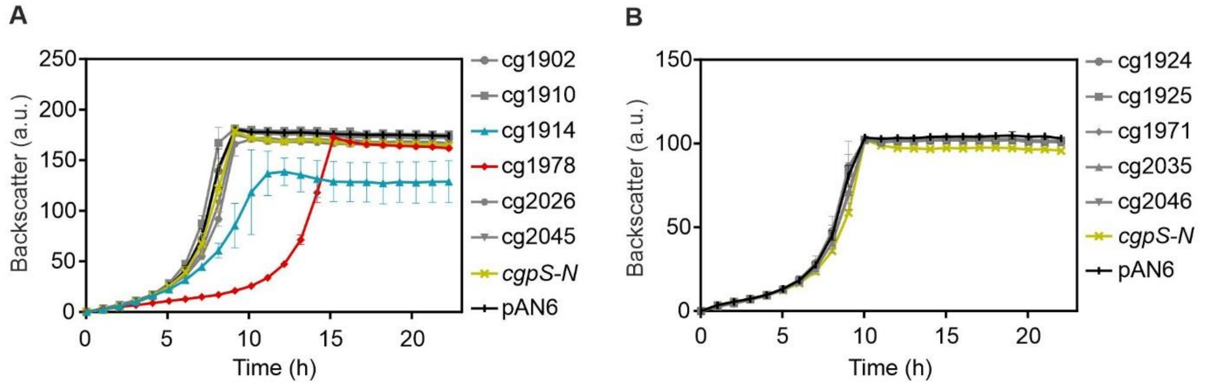


**Figure S1: Screening of small phagic proteins regarding their impact on cellular growth and CGP3 induction in *C. glutamicum*.** The cultivation of *C. glutamicum* ATCC 13032::$P_{lys}$-*eyfp* strains carrying the corresponding gene sequences of the small proteins on the pAN6 vector (under control of $P_{tac}$) was performed in CGXII medium with 2 % (w/v) glucose and 50 µM IPTG for 24 h. **(A,B)** Growth curves upon small protein overproduction are based on the backscatter measurements in the BioLector® microcultivation system. All data represent mean values with standard deviations from three independent biological triplicates (n=3). As the backscatter values were measured with two different BioLector® devices, data are represented in two separated graphs.

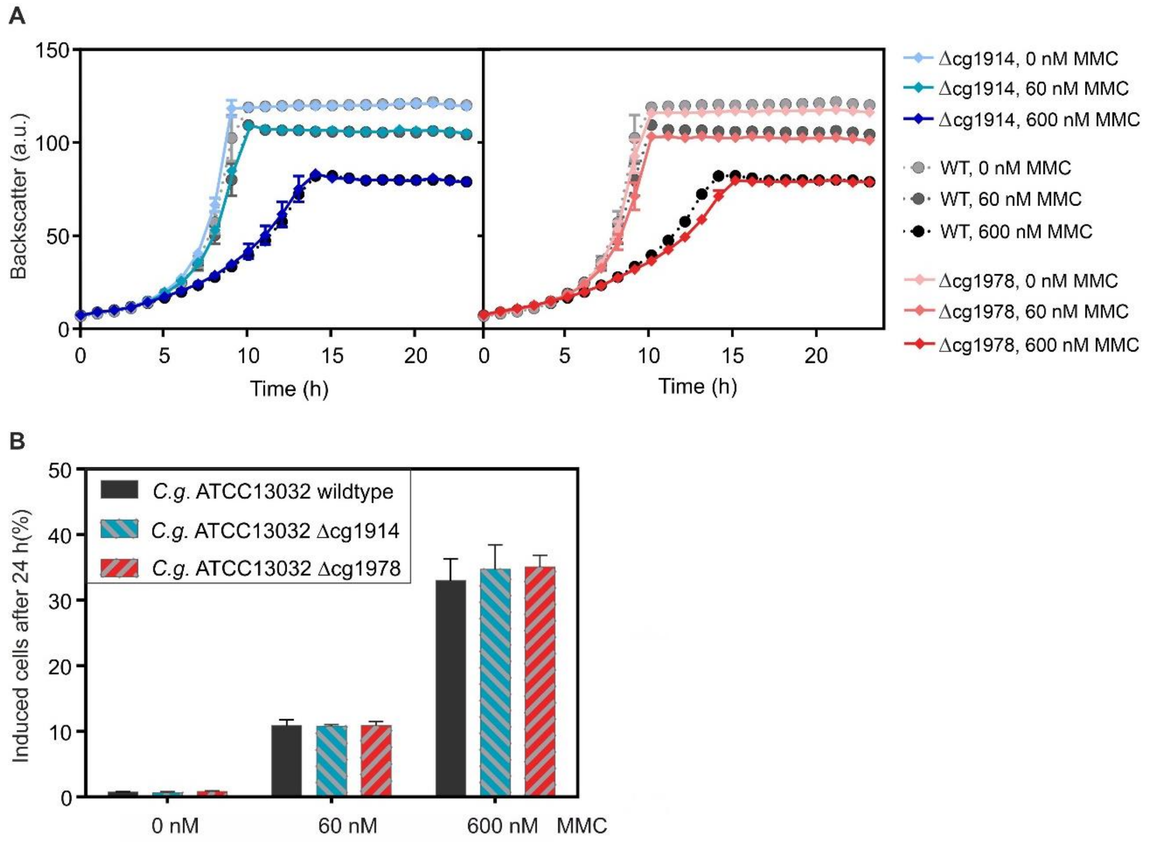


**Figure S2: Cell growth and prophage inducibility in the Δcg1914 or Δcg1978 mutant strain.** Cultivation of the *C. glutamicum* ATCC 13032 Δcg1914 strain, the *C. glutamicum* ATCC 13032 Δcg1978 strain and the *C. glutamicum* ATCC 13032 wildtype strain carrying the plasmid-based prophage reporter pJC1-$P_{lys}$-*eyfp* was performed in the BioLector® microcultivation system in CGXII medium with 2 % (w/v) glucose. All data represent mean values with standard deviations from three independent biological triplicates (n=3). **(A)** Growth curves based on the backscatter measurements in the BioLector® microcultivation system. **(B)** Percentage of induced cells after 24 h cultivation based on the flow cytometric measurements of the plasmid-based prophage reporter.

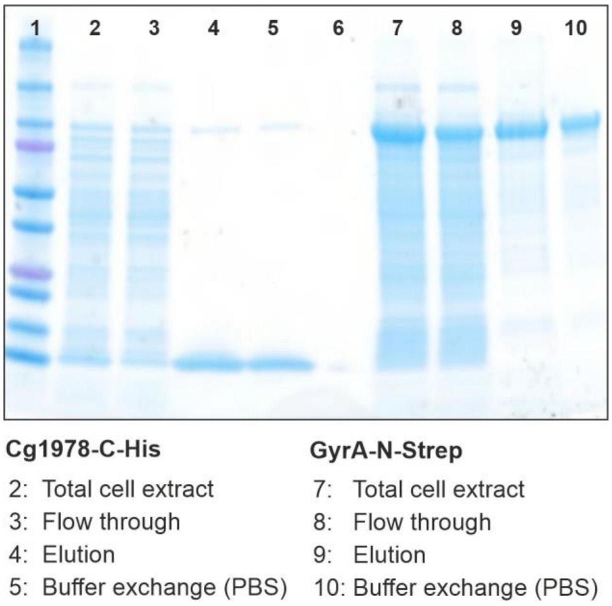


**Figure S3: Purification of proteins for surface plasmon resonance analysis of protein-protein interaction.** Cg1978 containing a C-terminal His-tag and GyrA containing a N-terminal Strep-tag were overproduced in *E. coli* BL21 (DE3) and purified via affinity purification. The gel electrophoresis was performed with 4-20 % gradient gels at 120 V for 60 min using the Precision Plus Protein™ Dual Xtra protein marker as a standard.

**Sequence alignment GyrA (E: *E. coli*, C: *C. glutamicum*, M: *M. tuberculosis*)**

```
E.    1 -------MSDLAREITPVNIEEELKSSYLDYAMSVIVGRALPDVRDGLKPVHRRVLYAMN
C.    1 ---MSDDNTGQFDRVNPIDINEEMQSSYIDYAMSVIVGRALPEVRDGLKPVHRRVLYAMF
M.    1 MTDTTLPPDDSLDRIEPVDIQQEMQRSYIDYAMSVIVGRALPEVRDGLKPVHRRVLYAMF

E.   54 VLGNDWNKAYKKSARVVGDVIGKYHPHGDSAVYDTIVRMAQPFSLRYMLVDGQGNFGSID
C.   58 DNGYRPDRSYVKSAKPVADTMGNFHPHGDTAIYDTLVRMAQPWSMRYPLVDGQGNFGSRG
M.   61 DSGFRPDRSHAKSARSVAETMGNYHPHGDASIYDTLVRMAQPWSLRYPLVDGQGNFGSPG

E.  114 GDSAAAMRYTEIRLAKIAHELMADLEKETVDFVDNYDGTEKIPDVMPTKIPNLLVNGSSG
C.  118 NDGPAAMRYTECRMTPLAMEMVRDIRENTVNFSPNYDGKTLEPDVLPSRVPNLLMNGSGG
M.  121 NDPPAAMRYTEARLTPLAMEMLREIDEETVDFIPNYDGRVQEPTVLPSRFPNLLANGSGG

E.  174 IAVGMATNIPPHNLTEVINGCLAYIDD----EDISIEGLMEHIPGPDFPTAAIINGRRGI
C.  178 IAVGMATNIPPHNLNELADAIFWLLENPDAEESEALEACMKFVKGPDFPTAGLIIGDKGI
M.  181 IAVGMATNIPPHNLRELADAVFWALENHDADEEETLAAVMGRVKGPDFPTAGLIVGSQGT

E.  230 EEAYRTGRGKVYIRARAEVEVDAKTGRETIIVHEIPYQVNKARLIEKIAELVKEKRVEGI
C.  238 HDAYTTGRGSIRMRGVTSIEEEG--NRTVIVITELPYQVNPDNLISNIAEQVRDGKLVGI
M.  241 ADAYKTGRGSIRMRGVVEVEEDS-RGRTSLVITELPYQVNHDNFITSIAEQVRDGKLAGI

E.  290 SALRDE-SDKDGMRIVIEVKRDAVGEVVLNNLYSQTQLQVSFGINMVALHHGQPKIMNLK
C.  296 SKIEDESSDRVGMRIVVTLKRDAVARVVLNNLFKHSQLQANFGANMLSIVDGVPRTLRLD
M.  300 SNIEDQSSDRVGLRIVIEIKRDAVAKVVINNLYKHTQLQTSFGANMLAIVDGVPRTLRLD

E.  349 DIIAAFVRHRREVVTRRTIFELRKARDRAHILEALAVALANIDPIIELIRHAPTPAEAKT
C.  356 QMLRYYVAHQIEVIVRRTQYRLDKAEERAHLLRGLVKALDMLDEVIALIRRSPTPDEART
M.  360 QLIRYYVDHQLDVIVRRTTYRLRKANERAHILRGLVKALDALDEVIALIRASETVDIARA

E.  409 ALVANPWQLGNVAAMLERAGDDAARPEWLEPEFGVRDGLYYLTEQQAQAILDLRLQKLTG
C.  416 GLMS----------------------------------LLDVDEAQADAILAMQLRRLAA
M.  420 GLIE----------------------------------LLDIDEIQAQAILDMQLRRLAA

E.  469 LEHEKLLDEYKELLDQIAELLRILGSADRLMEVIREELELVREQFGDKRRTEITANSADI
C.  442 LERQKIIDELAEIELEIADLKAILASPERQRTIVRDELTEIVEKYGDERRSQIIAATGDV
M.  446 LERQRIIDDLAKIEAEIADLEDILAKPERQRGIVRDELAEIVDRHGDDRRTRIIAADGDV

E.  529 NLEDLITQEDVVVTLSHQGYVKYQPLSEYEAQRRGGKGKSAARIKEEDFIDRLLVANTHD
C.  502 SEEDLIARENVVITITSTGYAKRTKVDAYKSQKRGGKGVRGAELKQDDIVRHFFVSSTHD
M.  506 SDEDLIAREDVVVTITETGYAKRTKTDLYRSQKRGGKGVQGAGLKQDDIVAHFFVCSTHD

E.  589 HILCFSSRGRVYSMKVYQLPEATRGARGRPIVNLLPLEQDERITAILPVTEFEEGVKVFM
C.  562 WILFFTNYGRVYRLKAFELPEASRTARGQHVANLLEFQPGEQIAQVIQLESYNDFPYLVL
M.  566 LILFFTTQGRVYRAKAYDLPEASRTARGQHVANLLAFQPEERIAQVIQIRGYTDAPYLVL

E.  649 ATANGTVKKTVLTEFNRLRTAGKVAIKLVDGDELIGVDLTSGEDEVMLFSAEGKVVRFKE
C.  622 ATAHGRVKKSRLLDYESARSGGLIAINLNEDDRLIGAALCGEEDDLLLVSEFGQSIRFTA
M.  626 ATRNGLVKKSKLTDFDSNRSGGIVAVNLRDNDELVGAVLCSADDDLLLVSANGQSIRFSA
```

```
E.   709 --SSVRAMGCNTTGVRGIRLGEGDKVVSLIVPRGDGAILTATQNGYGKRTAVAEYPTKSR
C.   682 DDEQLRPMGRATAGVKGMRFRDNDQLLSMSVVRDGEFLLVATSGGYGKRTPLEDYSTQGR
M.   686 TDEALRPMGRATSGVQGMRFNIDDRLLSLNVVREGTYLLVATSGGYAKRTAIEEYPVQGR

E.   767 ATKGVISIKVTERNGLVVGAVQVDDCDQIMMITDAGTLVRTRVSEISIVGRNTQGVILIR
C.   742 GGLGVVTFKYTPKRGRLVSAIAVEEDDEIFAITSAGGVVRTEVKQIRPSSRATMGVRLVN
M.   746 GGKGVLTVMYDRRRGRLVGALIVDDDSELYAVTSGGGVIRTAARQVRKAGRQTKGVRLMN

E.   827 TAEDENVVGLQRVAEPVDEEDLDTIDGSAAEGDDEIAPEV--D-VDDEPEE---E
C.   802 LEEGVELLAIDKNVEDQGEASAEAVAKGAVEGPASKTAAEETDSVDNGSDENGEE
M.   806 LGEGDTLLAIARNAEESGDDNAVDANGADQTGN----------------------
```

**Figure S4: Sequence alignment of the DNA gyrase subunit A from *C. glutamicum* (C), *M. tuberculosis* (M) and *E. coli* (E).** The multiple sequence alignment was conducted by using the Clustal Omega platform (Sievers et al., 2011). The output file was further transformed using Boxshade (http://www.ch.embnet.org/software/BOX_form.html). The sequence similarity of the DNA gyrase subunit A between *C. glutamicum* and *M. tuberculosis* is 71.46 %, the one between *C. glutamicum* and *E. coli* is 45.17 %.

**Sequence alignment GyrB (E: *E. coli*, C: *C. glutamicum*, M: *M. tuberculosis*)**

```
E.    1 ------------------------------MSNSYDSSSIKVLKGLDAVRKRPGMYIGDT
C.    1 MRGTTWGPKRVRWKRLYRISSEECSLKVANTEHNYDASSITILEGLEAVRKRPGMYIGST
M.    1 -----------------------MAAQKKKAQDEYGAASITILEGLEAVRKRPGMYIGST

E.   31 DDGTGLHHMVFEVVDNAIDEALAGHCKEIIVTIHADNSVSVQDDGRGIPTGIHPEEGVSA
C.   61 G-PRGLHHLIWEVVDNSVDEAMAGHATKVEVTLLEDGGVQVVDDGRGIPVDMHPS-GAPT
M.   38 G-ERGLHHLIWEVVDNAVDEAMAGYATTVNVVLLEDGGVEVADDGRGIPVATHAS-GIPT

E.   91 AEVIMTVLHAGGKFDDNSYKVSGGLHGVGVSVVNALSQKLELVIQREGKIHRQIYEHGVP
C.  119 VQVVMTQLHAGGKFDSDSYAVSGGLHGVGISVVNALSTRVEADIKLHGKHWYQNFEKSVP
M.   96 VDVVMTQLHAGGKFDSDAYAISGGLHGVGVSVVNALSTRLEVEIKRDGYEWSQVYEKSEP

E.  151 QAPLAVTGETEKTGTMVRFWPSLETFTNVTEFEYEILAKRLRELSFLNSGVSIRLRDKRD
C.  179 DE-LIEGGNARGTGTTIRFWPDAEIFE-TTEFDFETISRRLQEMAFLNKGLTITLTDNRA
M.  156 LG-LKQGAPTKKTGSTVRFWADPAVFE-TTEYDFETVARRLQEMAFLNKGLTINLTDERV

E.  211 ---------------------------------------------GKEDHFHYEGGIKAF
C.  237 TDEELELEALAEQGETATELSLDEIDNETELVEETTDAPKKPKKREKKKIFHYPNGLEDY
M.  214 TQDEVVDEVVSDVAEAPKSAS------------E-RAAESTAPHKVKSRTFHYPGGLVDF

E.  226 VEYLNKNKTPIHPNIFYFSTEKDGIGVEVALQWNDGFQENIYCFTNNIPQRDGGTHLAGF
C.  297 VHYLNRSKTNIHPSIVSFEAKGDDHEVEVAMQWNSSYKESVHTFANTINTREGGTHEEGF
M.  261 VKHINRTKNAIHSSIVDFSGKGTGHEVEIAMQWNAGYSESVHTFANTINTHEGGTHEEGF

E.  286 RAAMTRTLNAYMDKEGYSKKAKVSATGDDAREGLIAVVSVKVPDPKFSSQTKDKLVSSEV
C.  357 RSALTSLMNRYAREHKLLKEKEANLTGDDCREGLSAVISVRVGDPQFEGQTKTKLGNTEI
M.  321 RSALTSVVNKYAKDRKLLKDKDPNLTGDDIREGLAAVISVKVSEPQFEGQTKTKLGNTEV

E.  346 KSAVEQQMNELLAEYLLENPTDAKIVVGKIIDAARAREAARRAREMTRRKGALDLAGLPG
C.  417 KSFVQRMANEHIGHWLEANPAEAKVIINKAVGSAQARLAARKARDLVRRKSATDLGGLPG
M.  381 KSFVQKVCNEQLTHWFEANPTDAKVVVNKAVSSAQARIAARKARELVRRKSATDIGGLPG

E.  406 KLADCQERDPALSELYLVEGDSAGGSAKQGRNRKNQAILPLKGKILNVEKARFDKMLSSQ
C.  477 KLADCRSKDPEKSELYIVEGDSAGGSAKSGRDSMFQAILPLRGKILNVEKARLDKVLKNA
M.  441 KLADCRSTDPRKSELYVVEGDSAGGSAKSGRDSMFQAILPLRGKIINVEKARIDRVLKNT

E.  466 EVATLITALGCGIGRDEYNPDKLRYHSIIIMTDADVDGSHIRTLLLTFFYRQMPEIVERG
C.  537 EVQAIITALGTGIH-DEFDINKLRYHKIVLMADADVDGQHIATLLLTLLFRFMPDLVAEG
M.  501 EVQAIITALGTGIH-DEFDIGKLRYHKIVLMADADVDGQHISTLLLTLLFRFMRPLIENG

E.  526 HVYIAQPPLYKVKKGKQEQYIKDDEAMDQYQISIALDGATLHTNASAPALAGEALEKLVS
C.  596 HVYLAQPPLYKLKWQRGEPGFAYS------------------------------------
M.  560 HVFLAQPPLYKLKWQRSDPEFAYS------------------------------------

E.  586 EYNATQKMINRMERRYPKAMLKELIYQPTLTEADLSDEQTVTRWVNALVSELNDKEQHGS
C.  620 ------------------------------------DE-----------------ERD--
M.  584 ------------------------------------DR-----------------ERD--
```

```
E.   646 QWKFDVHTNAEQNLFEPIVRVRTHGVDTDYPLDHEFITGGEYRRICTLGEKLRGLLEEDA
C.   625 -----------------------------------------------------EQLN---
M.   589 -----------------------------------------------------GLLE---

E.   706 FIERGERRQPVASFEQALDWLVKESRRGLSIQRYKGLGEMNPEQLWETTMDPESRRMLRV
C.   629 ---EGLA-------------AGRKINKDDGIQRYKGLGEMNASELWETTMDPTVRILRRV
M.   593 ---AGLK-------------AGKKINKEDGIQRYKGLGEMDAKELWETTMDPSVRVLRQV

E.   766 TVKDAIAADQLFTTLMGDAVEPRRAFIEENALKAANIDI
C.   673 DITDAQRADELFSILMGDDVVARRSFITRNAKDVRFLDI
M.   637 TLDDAAAADELFSILMGEDVDARRSFITRNAKDVRFLDV
```

**Figure S5: Sequence alignment of the DNA gyrase subunit B from *C. glutamicum* (C), *M. tuberculosis* (M) and *E. coli* (E).** The multiple sequence alignment was conducted by using the Clustal Omega platform (Sievers et al., 2011). The output file was further transformed using Boxshade (http://www.ch.embnet.org/software/BOX_form.html). The sequence similarity of the DNA gyrase subunit B between *C. glutamicum* and *M. tuberculosis* is 73.48 %, the one between *C. glutamicum* and *E. coli* is 53.77 %.

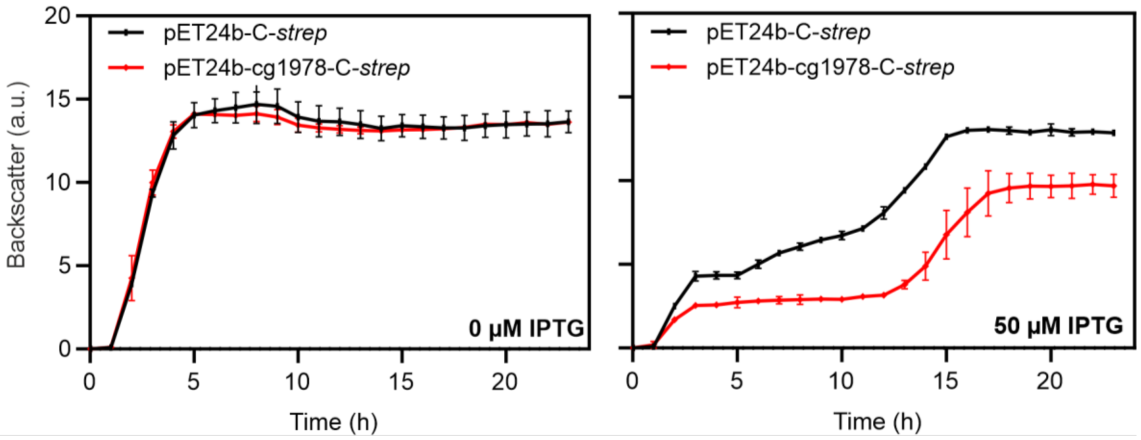


**Figure S6: Cell growth of *E. coli* BL21 (DE3) upon Cg1978 overproduction.** Cultivation of the *E. coli* BL21 (DE3) carrying the overexpression plasmid pET24b-cg1978-C-*strep* as well as the empty plasmid pET24b-C-*strep* was performed in the BioLector® microcultivation. For that, single colonies were used to inoculate pre-cultures in LB-$Kan_{50}$ medium. After cultivation overnight at 37°c and 120 rpm, the pre-cultures were used to inoculate main cultures in LB-$Kan_{50}$ medium to an $OD_{600}$ = 0.2. Cultivation was conducted at 37 °C and 1200 rpm and gene expression was induced by addition of 50 µM IPTG. During cultivation, biomass was measured as a function of backscattered light intensity with an excitation wavelength of 620 nm (filter module: λEx/ λEm: 620 nm/ 620 nm, gain: 15). All data represent mean values with standard deviations from three independent biological triplicates (n=3).

**Table S1: Bacterial strains used in this study**

| Strain | Genotype and relevant characteristics | Reference |
|---|---|---|
| *E. coli* BL21 (DE3) | $F^-$ *ompT hsdS*$_B$(*r*$_B^-$ *m*$_B^-$) *gal dcm* λ(DE3) | (Studier & Moffatt, 1986) |
| *E. coli* DH5α | *supE44 ΔlacU169 (f80lacZ*DM15*) hsdR17 recA1 endA1 gyrA96 thi-1 relA1* | Invitrogen |
| *C. glutamicum* ATCC 13032 | Biotin-auxotrophic wild type | (Ikeda & Nakagawa, 2003) |
| ATCC 13032::P$_{lys}$-*eyfp* | ATCC 13032 with promoter fusion P$_{lys}$-*eyfp* integrated into the intergenic region of cg1121 and cg1122 | (Helfrich et al., 2015) |
| ATCC 13032 Δ*recA*::P$_{lys}$-*eyfp* | ATCC 13032::P$_{lys}$-*eyfp* with in-frame deletion of ATPase domain of *recA* (cg2141) | (Helfrich et al., 2015) |
| ATCC 13032::P$_{recA}$-*venus* | ATCC 13032 with promoter fusion P$_{recA}$-eyfp integrated into the intergenic region ofcg1121 and cg1122 | (Helfrich et al., 2015) |
| MB001 | ATCC 13032 ΔCGP1 (cg1507-cg1524), ΔCGP2 (cg1746-cg1752) and ΔCGP3 (cg1890-cg2071) (BA strain) | (Baumgart, Unthan, et al., 2013) |
| ATCC 13032 Δcg1914 | ATCC 13032 with in-frame deletion of cg1914 | This work |
| ATCC 13032 Δcg1978 | ATCC 13032 with in-frame deletion of cg1978 | This work |

**Table S2: Plasmids used in this study.** Numbers represent oligonucleotides used for amplification of the insert DNA (Table S3) using the *C. glutamicum* genome as template. The vectors were linearized with the indicated restriction enzyme and plasmids were constructed using Gibson assembly. Sequencing was performed using the listed oligonucleotides.

| Plasmids | Template | Primer | Vector | Restriction enzymes | Sequencing primer |
|---|---|---|---|---|---|
| **Plasmids for overproduction of small proteins in *C. glutamicum*** | | | | | |
| pAN6 | Kan$^R$; *C. glutamicum*/ *E. coli* shuttle vector for regulated gene expression; derivative of pEKEx2 ($P_{tac}$, *lacI*$^q$, pBL1 $oriV_{C.g.}$, pUC18 $oriV_{E.c.}$) (Frunzke et al., 2008) | | | | |
| pAN6-*cgpS-N* | Kan$^R$; pAN6 derivative containing a truncated variant of the *cgpS* gene under control of $P_{tac}$ (Pfeifer et al., 2016) | | | | |
| pAN6-cg1902 | *C. glutamicum* chromosome | 1 + 2 | pAN6 | NdeI + EcoRI | 45 + 46 |
| pAN6-cg1910 | *C. glutamicum* chromosome | 3 + 4 | pAN6 | NdeI + EcoRI | 45 + 46 |
| pAN6-cg1914 | *C. glutamicum* chromosome | 5 + 6 | pAN6 | NdeI + EcoRI | 45 + 46 |
| pAN6-cg1924 | *C. glutamicum* chromosome | 7 + 8 | pAN6 | NdeI + EcoRI | 45 + 46 |
| pAN6-cg1925 | *C. glutamicum* chromosome | 9 + 10 | pAN6 | NdeI + EcoRI | 45 + 46 |
| pAN6-cg1971 | *C. glutamicum* chromosome | 11 + 12 | pAN6 | NdeI + EcoRI | 45 + 46 |
| pAN6-cg1978 / pAN6-*gip* | *C. glutamicum* chromosome | 13 + 14 | pAN6 | NdeI + EcoRI | 45 + 46 |
| pAN6-cg2026 | *C. glutamicum* chromosome | 15 + 16 | pAN6 | NdeI + EcoRI | 45 + 46 |
| pAN6-cg2035 | *C. glutamicum* chromosome | 17 + 18 | pAN6 | NdeI + EcoRI | 45 + 46 |
| pAN6-cg2045 | *C. glutamicum* chromosome | 19 + 20 | pAN6 | NdeI + EcoRI | 45 + 46 |
| pAN6-cg2046 | *C. glutamicum* chromosome | 21 + 22 | pAN6 | NdeI + EcoRI | 45 + 46 |
| **Plasmids for overproduction of small proteins with affinity tags in *E. coli*** | | | | | |
| pET24b | Kan$^R$; *E. coli* vector for regulated gene expression; derivative of pBR322 ($P_{T7}$, *lacI*$^q$, f1 ori, pBR322 ori, T7 terminator) (Novagen) | | | | |
| pET24b-cg1978-C-*strep* | *C. glutamicum* chromosome | 23 + 24 | pET24b | NdeI + NheI | 47 + 48 |
| pET24b-cg1978-C-*his* | *C. glutamicum* chromosome | 25 + 26 | pET24b | NdeI + NheI | 47 + 48 |
| pET24b-*gyrA*-N-*strep* | *C. glutamicum* chromosome | 27 + 28 | pET24b | BlpI + NheI | 47 + 48<br>49 + 50 |

| | | | | | |
|---|---|---|---|---|---|
| pET24b-*gyrB*-C-*strep* | *C. glutamicum* chromosome | 29 + 30 | pET24b | NdeI + NheI | 47 + 48<br>51 + 52 |
| **Plasmids for construction of deletion mutants** | | | | | |
| pK19*mobsacB* | Plasmid containing a negative (*sacB*) as well as a positive selection marker (*kan*$^R$) for allelic exchange in *C. glutamicum* (pK18 oriV$_{E.c}$, *sacB, lacZ*α) (Schäfer et al., 1994) | | | | |
| pK19*mobsacB*-Δcg1914 | *C. glutamicum* chromosome | 31 + 32<br>33 + 34 | pK19*mobsacB* | EcoRI + HindIII | 53 + 54 |
| pK19*mobsacB*-Δcg1978 | *C. glutamicum* chromosome | 35 + 36<br>37 + 38 | pK19*mobsacB* | EcoRI + HindIII | 53 + 54 |
| **Reporter plasmids** | | | | | |
| pJC1 | Kan$^R$, Amp$^R$, *C. glutamicum* shuttle vector (Cremer et al., 1990) | | | | |
| pJC1-*venus*-term | pJC1 derivative carrying the venus coding sequence and additional terminators (Baumgart, Luder, et al., 2013) | | | | |
| pJC1-P$_{lys}$-*venus* | *C. glutamicum* chromosom | 39 + 40 | pJC1-*venus*-term | BamHI + NdeI | 55 + 56 |

### Table S3: Oligonucleotides used in this study

| Primer No. | Oligonucleotide name | Sequence (5' → 3') |
|---|---|---|
| **Construction of plasmids for overproduction of small proteins in *C. glutamicum*** | | |
| 1 | pAN6_cg1902_fw | TACATATGACCTGAGCTAGCATGATTAAGAGACTGGCTGCAGG |
| 2 | pAN6_cg1902_rv | AAACGACGGCCAGTGAATTCTTATACAACTTGAATAGCCGTACCTG |
| 3 | pAN6_cg1910_fw | TACATATGACCTGAGCTAGCTTGGAGTTTATTTTATTTTTTGCACTT |
| 4 | pAN6_cg1910_rv | AAACGACGGCCAGTGAATTCCTATTTTTCCAACTTGTCGCTCTTAC |
| 5 | pAN6_cg1914_fw | TACATATGACCTGAGCTAGCATGAACTGCCCAAACTGCTC |
| 6 | pAN6_cg1914_rv | AAACGACGGCCAGTGAATTCTTACAGTACTTCGATATATCCGCAGTC |
| 7 | pAN6_cg1924_fw | CTGCAGAAGGAGATATACATATGTACTCGACATCATCATTTACC |
| 8 | pAN6_cg1924_rv | AAACGACGGCCAGTGAATTCTTACTGCTCATTATGAGGTGCC |
| 9 | pAN6_cg1925_fw | CTGCAGAAGGAGATATACATATGGTGGTGTGCGGC |
| 10 | pAN6_cg1925_rv | AAACGACGGCCAGTGAATTCTTACGGCTGTCGAGCTG |
| 11 | pAN6_cg1971_fw | CTGCAGAAGGAGATATACATATGTCTAATCTCGGCACATACTATG |
| 12 | pAN6_cg1971_rv | AAACGACGGCCAGTGAATTCTCAGAAACCAGGCTGTTGAGAC |
| 13 | pAN6_cg1978_fw | TACATATGACCTGAGCTAGCATGGCTAAAGAATTCGAATTCACC |
| 14 | pAN6_cg1978_rv | AAACGACGGCCAGTGAATTCTTACTCGACGATGACGTAGGG |
| 15 | pAN6_cg2026_fw | TACATATGACCTGAGCTAGCATGACCAAGCGAAATATCACTACTGT |
| 16 | pAN6_cg2026_rv | AAACGACGGCCAGTGAATTCTCAGCCCTTAGGTGGGTG |
| 17 | pAN6_cg2035_fw | CTGCAGAAGGAGATATACATATGTCAATCAATGCGTTCTGG |
| 18 | pAN6_cg2035_rv | AAACGACGGCCAGTGAATTCTCAGGCACCTAGATATGTGATTAC |
| 19 | pAN6_cg2045_fw | TACATATGACCTGAGCTAGCATGCCTCAACGCGAAAAGC |
| 20 | pAN6_cg2045_rv | AAACGACGGCCAGTGAATTCTTAGCTGTCCACAAGAATGCC |
| 21 | pAN6_cg2046_fw | CTGCAGAAGGAGATATACATATGGGGTACCTGGGAATTGATAG |
| 22 | pAN6_cg2046_rv | AAACGACGGCCAGTGAATTCTTAGATATCGACTCCTAGCGCTC |
| **Construction of plasmids for overproduction of proteins with affinity tags in *E. coli*** | | |
| 23 | cg1978_C-Strep_pET_fw | AAGAAGGAGATATACATATGATGGCTAAAGAATTCGAATTCACCATC |
| 24 | cg1978_C-Strep_pET_rv | AAATACAGGTTCTCGCTAGCCTCGACGATGACGTAGGG |
| 25 | cg1978_C-His_pET_fw | AAGAAGGAGATATACATATGATGGCTAAAGAATTCGAATTCACCATC |
| 26 | cg1978_C-His_pET_rv | AAATACAGGTTCTCGCTAGCCTCGACGATGACGTAGGG |
| 27 | *gyrA*_N-Strep_pET_fw | TGTATTTTCAGGGCGCTAGCGTGAGCGACGACAATACC |
| 28 | *gyrA*_N-Strep_pET_rv | TATGCTAGTTATTGCTCAGCTTATTCCTCGCCGTTTTCGTCG |
| 29 | *gyrB*_C-Strep_pET_fw | AAGAAGGAGATATACATATGGTGGCAAACACTGAACACAATTATGAC |
| 30 | *gyrB*_C-Strep_pET_rv | AAATACAGGTTCTCGCTAGCGATATCGAGGAAACGAACATCCTTGG |
| **Construction of plasmids for genomic deletion in *C. glutamicum*** | | |
| 31 | LF_fw_cg1914 | CCATGATTACGCCAAGCTTGTCTGTGGACATCATGAAAAACG |
| 32 | LF_rv_cg1914 | GTCTGTAACCGAGCATCTCTCGATGATCTCCTTTTAAGGGATTGAGGTG |
| 33 | RF_fw_cg1914 | GAGAGATGCTCGGTTACAGACCACGTCTGCAGCGACTG |
| 34 | RF_rv_cg1914 | AACGACGGCCAGTGAATTCGGACCATGAGCGGCCGGTT |
| 35 | LF_fw_cg1978 | ACCATGATTACGCCAAGCTTGACCAAATCGGCGATGTGTTTG |
| 36 | LF_rv_cg1978 | GTCTGTAACCGAGCATCTCTCAATAAGTGTTCTTTCTTATGCGAGGTG |
| 37 | RF_fw_cg1978 | GAGAGATGCTCGGTTACAGACAGGCGTTTCTTTTTCTCCCCC |

| | | |
|---|---|---|
| 38 | RF_rv_cg1978 | AAAACGACGGCCAGTGAATTGCCCGTGAGGTCACCCTTAT |

**Construction of prophage reporter plasmid**

| | | |
|---|---|---|
| 39 | pJC1_$P_{lys}$-*lys*_fw | GACGCCGCAGGGGGATCCCCTTCTTTGAGGCTTGATGCCTTGATC |
| 40 | pJC1_$P_{lys}$-*lys*_rv | CTCCTCGCCCTTGCTCACCATATGATATCTCCTTCTTAAAGTTCAATTTTTC GGCATTGCGCCTTTAATCG |

**Sequencing primer**

| | | |
|---|---|---|
| 41 | Δcg1914_fw_seq | AGGTGTGTATAACACCCGACAAG |
| 42 | Δcg1914_rv_seq | ATGCGACGCAAATTTTGGACC |
| 43 | Δcg1978_fw_seq | TCCGACATCATTCACATGACTGAC |
| 44 | Δcg1978_rv_seq | TCGTCAAGGTAGAGAGACATAAGTTATGTAG |
| 45 | pAN6_fw_seq | GATATGACCATGATTACGCCAAGC |
| 46 | pAN6_rv_seq | CGGCGTTTCACTTCTGAGTTCGGC |
| 47 | pET24b_fw_seq | CGATATAGGCGCCAGCAACC |
| 48 | pET24b_rv_seq | CCTCAAGACCCGTTTAGAGG |
| 49 | *gyrA*-N-strep _fw_seq | GGCTGACGCAATTCTGGCAATG |
| 50 | *gyrA*-N-strep _rv_seq | CAGCTCATCGCGAACGATGG |
| 51 | *gyrB*-C-strep _fw_seq | CGAGGAAGGTTTCCGCTCTG |
| 52 | *gyrB*-C-strep _rv_seq | GGAAATAACCGCGGACAGGC |
| 53 | pK19*mobSacB*_fw_seq | AGCGGATAACAATTTCACACAGGA |
| 54 | pK19*mobSacB*_rv_seq | CGCCAGGGTTTTCCCAGTCACGAC |
| 55 | pJC1-*venus*_fw_seq | TGAAGACCGTCAACCAAAGG |
| 56 | pJC-*venus*_rv_seq | CTGAACTTGTGGCCGTTTAC |

**Table S4: Impact of *gip* (cg1978) overexpression on global expression levels.** A genome-wide comparison of mRNA levels the *C. glutamicum* ATCC 13032 strain overexpressing *gip* and the wildtype strain carrying the empty vector control was performed. The shown mRNA ratios indicated mean values from three independent biological replicates (n=3). The strains were cultivated in CGXII medium with 2 % (w/v) glucose and mRNA was prepared from cells at an $OD_{600}$ of 6. The mRNA ratios were calculated by dividing the mRNA levels of the *gip* overexpressing strain by the mRNA levels of the strain carrying the empty vector control. The table includes all genes which showed a changed mRNA level in all experiments (mRNA ratio > 2.0: upregulation (red) or < 0.5: downregulation (green), p-value ≤ 0.05).

**The table is provided in a separate file (Table S4_Complete Microarray Data).**

**Video S1: Time lapse video of a *C. glutamicum* microcolony of the prophage reporter strain under cg1978 overexpression (50 µM IPTG).** Cells of the prophage reporter strain ATCC 13032::P$_{lys}$-*eyfp* carrying the overexpression plasmid pAN6-cg1978 were cultivated in microfluidic chambers (Grünberger et al., 2015) using CGXII minimal medium with 2% (w/v) glucose and 25 µg ml$^{-1}$ kanamycin for 18 h. Overexpression was induced using 50 µM IPTG.

**Video S2: Time lapse video of a *C. glutamicum* microcolony of the prophage reporter strain under standard conditions (0 µM IPTG).** The same reporter strain (Video S1) carrying the overexpression plasmid pAN6-cg1978 was grown in the absence of IPTG serving as a control for normal cell growth.

**The videos are provided as separate files:**

- Video S1_cg1978 overexpression_prophage reporter_50 µM IPTG
- Video S1_cg1978 overexpression_prophage reporter_0 µM IPTG
